# Amyloidogenesis of SARS-CoV-2 Spike Protein

**DOI:** 10.1101/2021.12.16.472920

**Authors:** Sofie Nyström, Per Hammarström

**Affiliations:** Dept of Physics, Chemistry and Biology, Linköping University, Linköping, Sweden

## Abstract

SARS-CoV-2 infection is associated with a surprising number of morbidities. Uncanny similarities with amyloid-disease associated blood coagulation and fibrinolytic disturbances together with neurologic and cardiac problems led us to investigate the amyloidogenicity of the SARS-CoV-2 Spike protein (S-protein). Amyloid fibril assays of peptide library mixtures and theoretical predictions identified seven amyloidogenic sequences within the S-protein. All seven peptides in isolation formed aggregates during incubation at 37°C. Three 20-amino acid long synthetic Spike peptides (sequence 191-210, 599-618, 1165-1184) fulfilled three amyloid fibril criteria: nucleation dependent polymerization kinetics by ThT, Congo red positivity and ultrastructural fibrillar morphology. Full-length folded S-protein did not form amyloid fibrils, but amyloid-like fibrils with evident branching were formed during 24 hours of S-protein co-incubation with the protease neutrophil elastase (NE) in vitro. NE efficiently cleaved S-protein rendering exposure of amyloidogenic segments and accumulation of the peptide 193-202, part of the most amyloidogenic synthetic Spike peptide. NE is overexpressed at inflamed sites of viral infection and at vaccine injection sites. Our data propose a molecular mechanism for amyloidogenesis of SARS-CoV-2 S-protein in humans facilitated by endoproteolysis. The potential implications of S-protein amyloidogenesis in COVID-19 disease associated pathogenesis and consequences following S-protein based vaccines should be addressed in understanding the disease, long COVID-19, and vaccine side effects.

---

The SARS-CoV-2 pandemic has severely impacted the world and official numbers state that over 270 million people have been infected (Dec 2021), but unrecorded cases are likely significantly higher. Corona viruses use the surface spike protein (S-protein) to attach to human cells. The S-protein is a homotrimer and each subunit of SARS-CoV-2 S-protein comprise 1273 amino acids. Four common corona viruses; OC43, 229E, NL63, HKU1 infect humans and colonize the respiratory tract. Recently emerged SARS, MERS and since 2019 also SARS-CoV-2, render severe disease. Although corona virus infections are common, it has not before COVID-19 been reports of such a wide distribution of complex symptoms involving other organs than the respiratory tract. Blood clotting, heart failure, peripheral neuropathy and CNS disorders are severe symptoms out of many reported. What could be the basis for this pathogenesis? Amyloidosis manifests as systemic and localized disorders with many overlapping phenotypes with reported COVID-19 symptoms. We therefore hypothesized on a potential molecular link. We were inspired by previous hypotheses about human and viral protein amyloids and interactions between them [1-3], in particular SARS-CoV spike proteins [4-6]. We asked the question if SARS-CoV-2 S-protein is amyloidogenic?

Motivated by Zhang et al 2018 [5] we first obtained a pool library of spike peptides, intended for antibody screening. The library comprised 316 peptides (delivered in two subpools of 158 peptides each) derived from a peptide scan (15-mers with 11 AA overlap, Supporting Inf. 2) through the entire SARS-CoV-2 S-protein (Protein ID: P0DTC2). The library was assayed for *in vitro* amyloid fibril formation. Fibrils were formed in both peptide subpools (Supporting Inf. 1 and SFig. 1). Encouraged by the results we moved on to generate 20 AA peptide sequences from the full-length SARS-CoV-2 S-protein. We aimed to address the most amyloidogenic sequences and used the WALTZ (https://waltz.switchlab.org/) prediction algorithm [7] to identify such segments (Table 1).

**Table 1.**
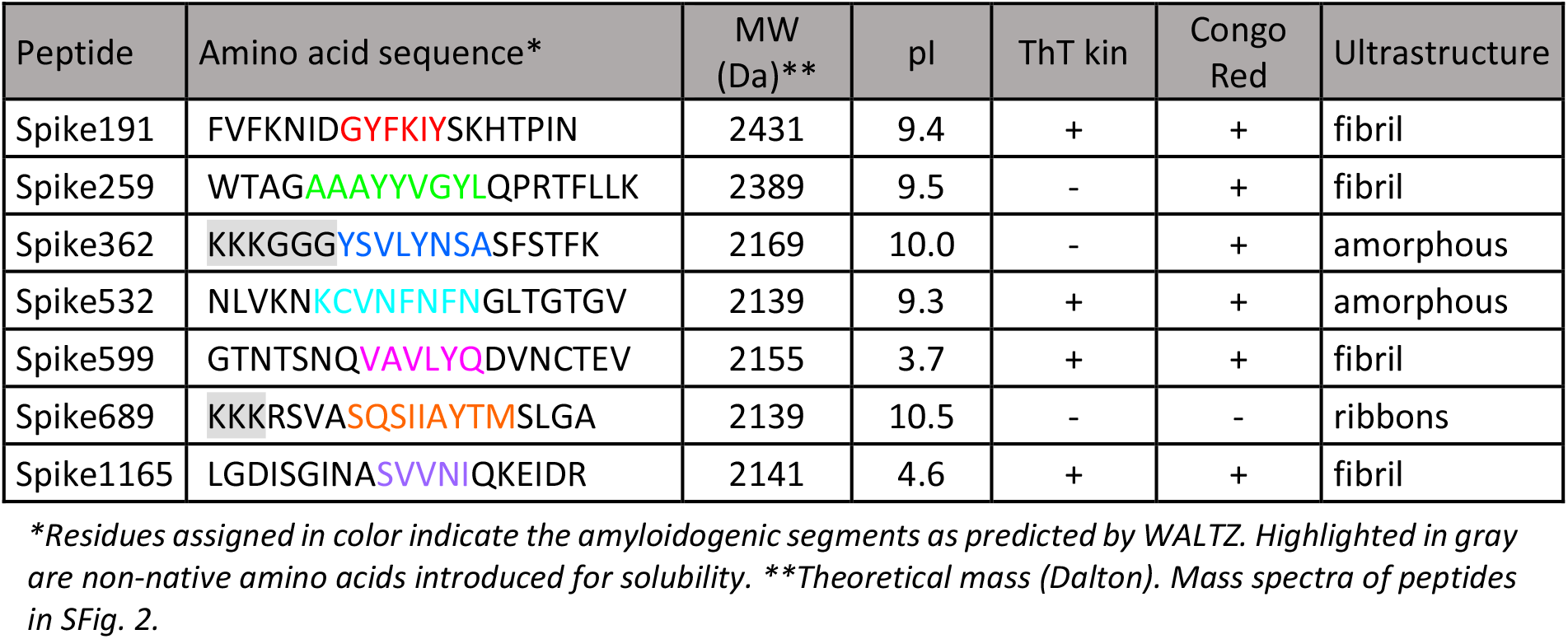
Amino acid sequences and properties of synthetic SARS-CoV-2 S-protein peptides.

The seven amyloidogenic sequences were distributed over the entire S-protein and are named according to the starting position of the S-protein (Fig. 1). All but one (Spike362) of the predicted sequences are in beta-sheet conformation in the cryo-EM structure of SARS-CoV-2 S in its closed state [8].

**Figure 1.**
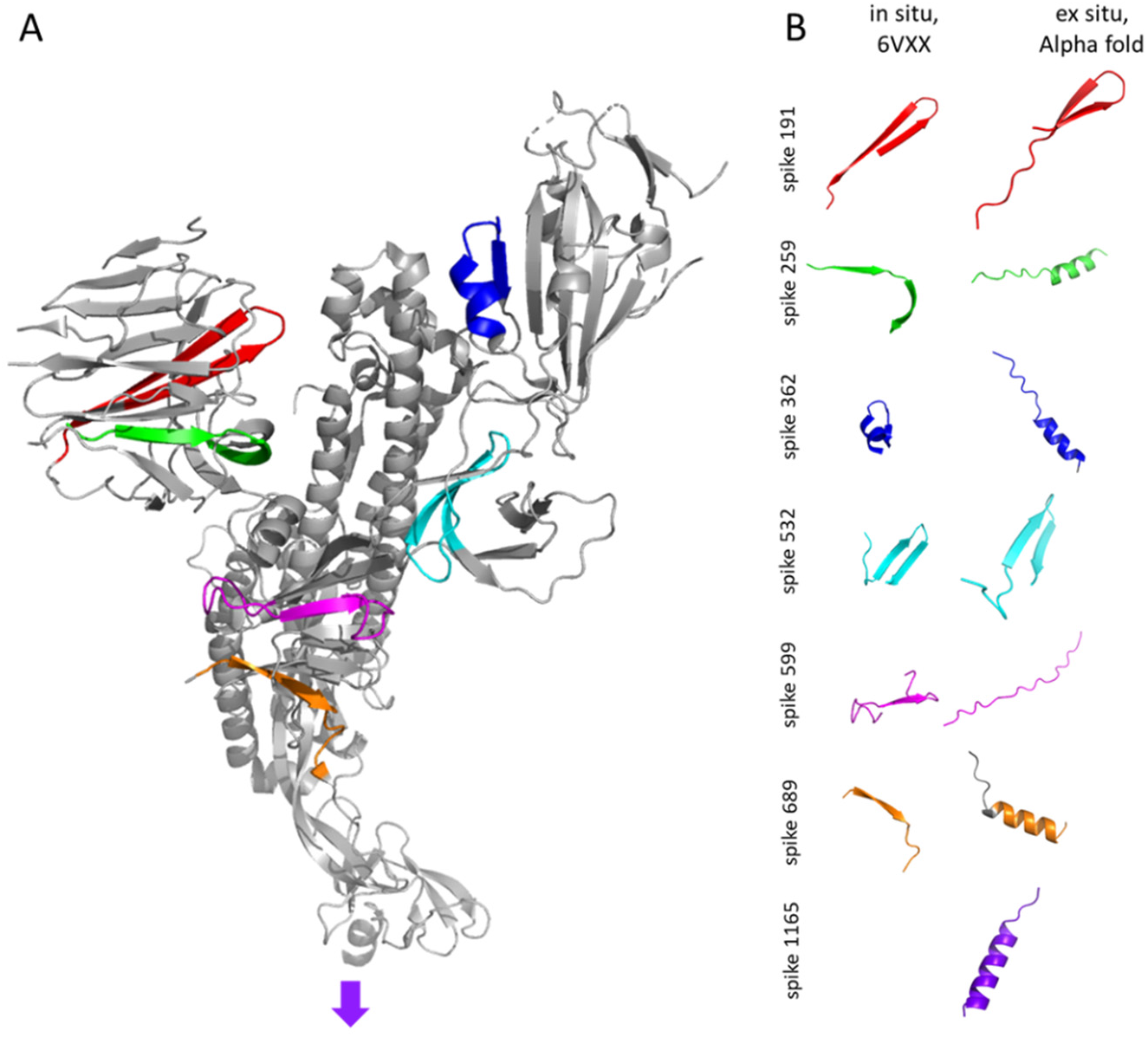
A. The structure of one protomer of the trimeric SARS-CoV-2 S-protein in its closed state, PDB code: 6VXX [8] with the predicted full sequence of the amyloidogenic peptides highlighted in the same colors as the predictions in Table 1. B. Conformation of peptides within the folded S-protein in comparison with Alpha Fold 2 models of the synthetic peptides (Table 1).

The membrane spanning C-terminal part of the protein (Spike1165) is not included in the structure. Results from Alpha fold 2 [9] on the selected peptides indicate that three peptides; Spike259, Spike362 and Spike689, that are in β-strand conformation in the full-length protein are in helical conformation as free peptides (Fig. 1B). One of the peptides (Spike599) was predicted by Alpha fold 2 to be random coil and the membrane spanning part was predicted to be an α-helix. Note that the sequences of Spike362 and Spike689 have been N-terminally modified with solubilizing sequences to enable production. Alpha fold 2 was performed on the sequences in Table 1 (Fig. 1B).

Lyophilized peptides were solubilized in hexa-fluoroisopropanol (HFIP) and were diluted in PBS-buffer (pH 7.5) to a final peptide concentration of 0.1 mg/ml (10% HFIP) and monitored for *in vitro* amyloid fibril formation kinetics using ThT, Congo red birefringence (CR) and negative stain transmission electron microscopy (TEM). Fibrils were formed within a few hours from most of the synthetic peptides by at least one but not all assays (Table 1, Fig. 2). Spike191, Spike532 and Spike1165 fulfilled all our amyloid criteria: sigmoidal ThT kinetics, Congophilicity and fibrillar ultrastructure (Fig. 2, Table 1). In particular, Spike191 formed exceptionally well-ordered fibrils (Fig. 2C, Fig. 3C).

**Figure 2.**
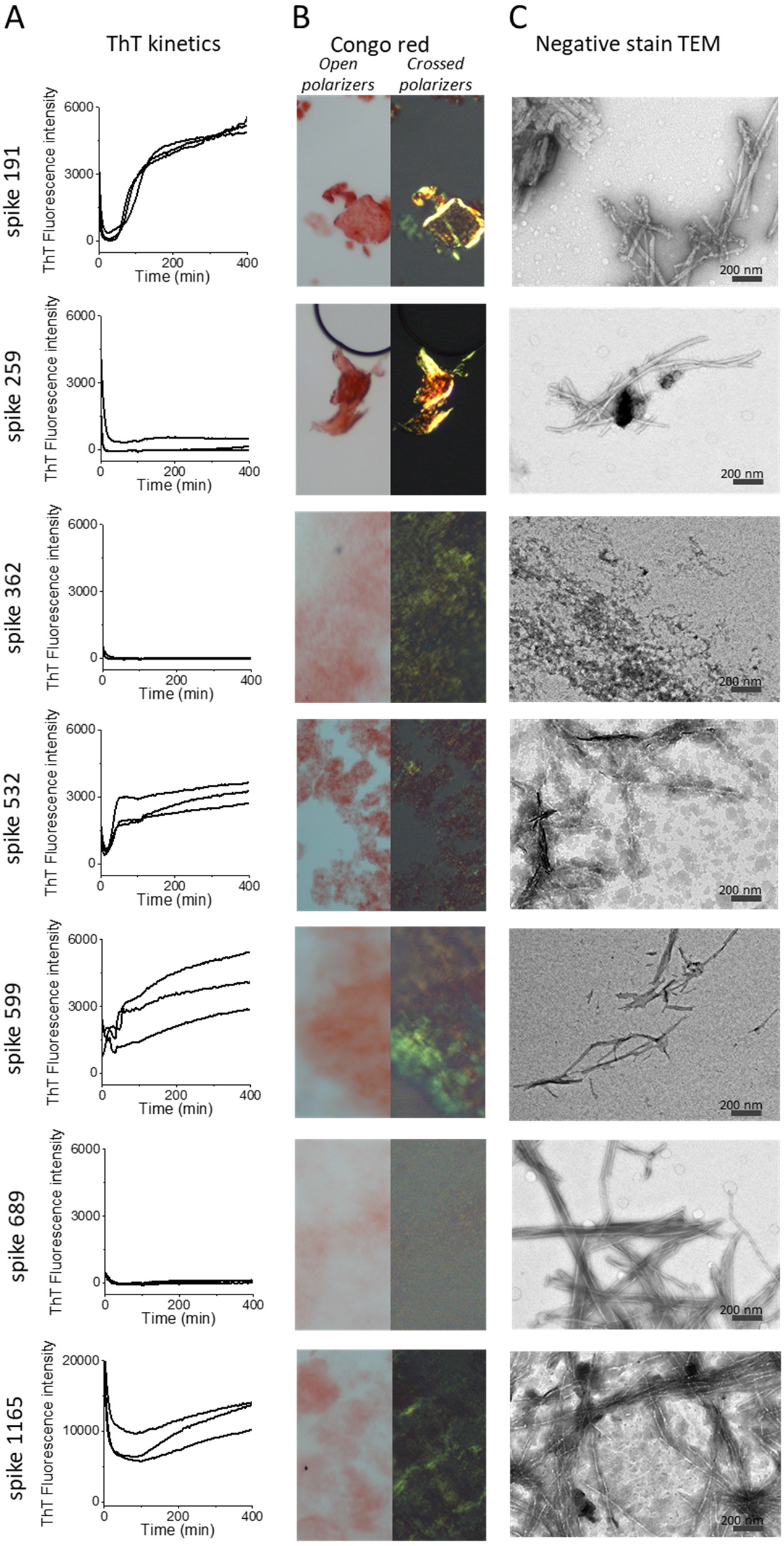
Amyloid fibril assays of SARS-CoV-2 S peptides (0.1 mg/ml). A. Fibril formation kinetics monitored by ThT fluorescence. B. Congo red staining microscopy with open and crossed polarizers. C. Ultrastructure by negative stain TEM.

**Figure 3.**
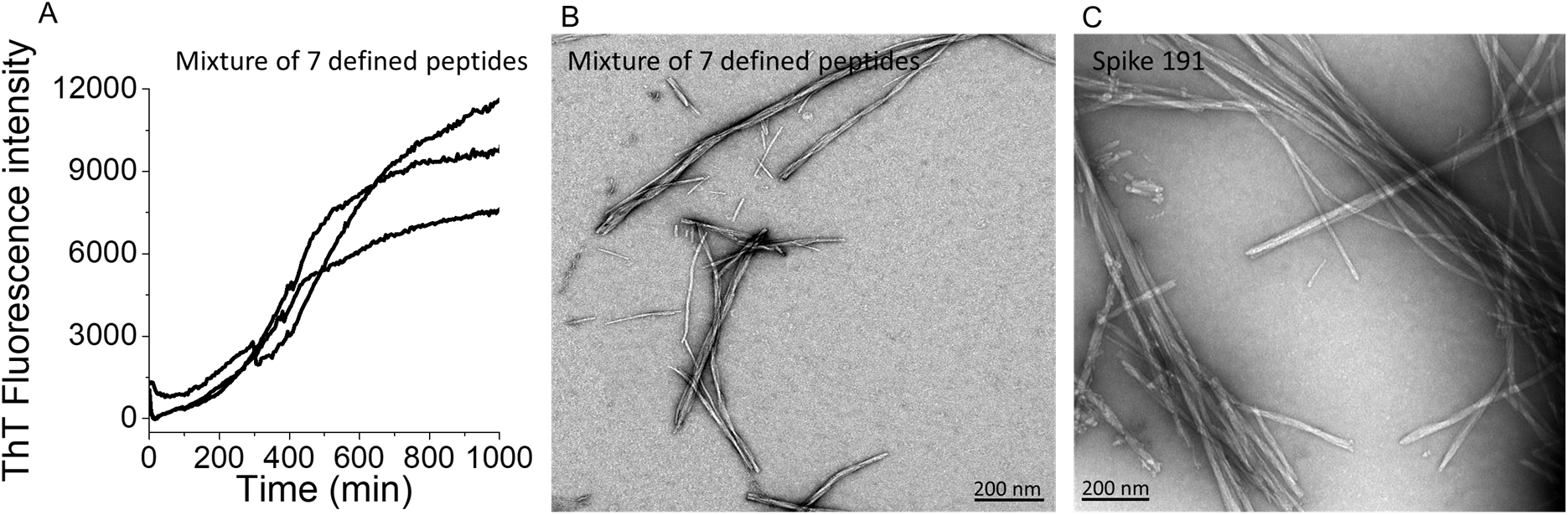
Amyloid fibrillation of 7 mixed SARS-CoV-2 S-peptides (total concentration 0.1 mg/ml). A. Fibril formation kinetics monitored by ThT fluorescence. B. Fibrillar structures by negative stain TEM. C. Fibrillar structures by negative stain TEM of Spike191 resembling the mix.

The synthetic peptides were also prepared as a mixture of all seven peptides with a final concentration of 0.1 mg/ml of total peptide (concentration of each peptide ∼0.014 mg/ml). The mixture of peptides resulted in amyloid fibril formation with sigmoidal ThT kinetics suggesting a nucleation dependent mechanism (Fig. 3A). The fibril morphology from the mixture (Fig. 3B) most closely resembled that of fibrils composed of the well-ordered Spike191 (cf. Fig 3B; Fig. 3C), suggesting that this peptide is dominating the fibril structures in the mixture

What would be a plausible mechanism for S-protein fibril formation in a SARS-CoV-2 infected patient? SARS-CoV-2 S-protein is fairly stable (T_m_ >50 °C) [10] (see below) and would not readily denature spontaneously. Also, such a large protein with complex fold will not easily misfold into an amyloid state. However, proteolysis is an obvious candidate mechanism.

Endoproteolysis of precursor proteins is a well-known molecular initiation mechanism in several amyloidoses notably Alzheimer’s disease (AβPP), British and Danish dementia (ABri/ADan), and Finnish familial amyloidosis (AGel). Proteolysis of the full-length protein is also evident in many other amyloid disease deposits from ATTR, ALys, AA, ASem1 [11, 12]. SARS-CoV-2 S-protein is proteolyzed during infection by host furin-like enzymes and by serine proteases such as the transmembrane protease, serine 2 (TMPRSS2), at the cell surface [13]. S-protein is further proteolyzed during inflammation. Neutrophils are the dominating class of leucocytes and are one of the first responders during acute inflammation. Neutrophils are recruited to the bronchoalveolar space of patients infected with a range of different respiratory viruses, including SARS-CoV-2 [14]. Neutrophils act both by phagocytosis of opsonized pathogens and by extracellular release of enzymes such as neutrophil elastase (NE). NE is a serine protease coupled to obstructive lung diseases such as cystic fibrosis, chronic obstructive pulmonary disease (COPD) [15] and alpha-1-antitrypsin deficiency [16]. The amino acid sequence of SARS-CoV-2 S-protein was tested with in silico proteolytic cleavage by NE using Expasy Peptide cutter. One of the resulting peptides, Spike193-212, was closely matching peptide Spike191, only frame shifted by 2 amino acids (Supporting information 3) implying a testable hypothesis.

We therefore subjected full-length SARS-CoV-2 S-protein to NE cleavage *in vitro*. We first determined that the S-protein was folded by thermal unfolding experiments by differential scanning fluorimetry (DSF) (Fig. 4, SFig. 3).

**Figure 4.**
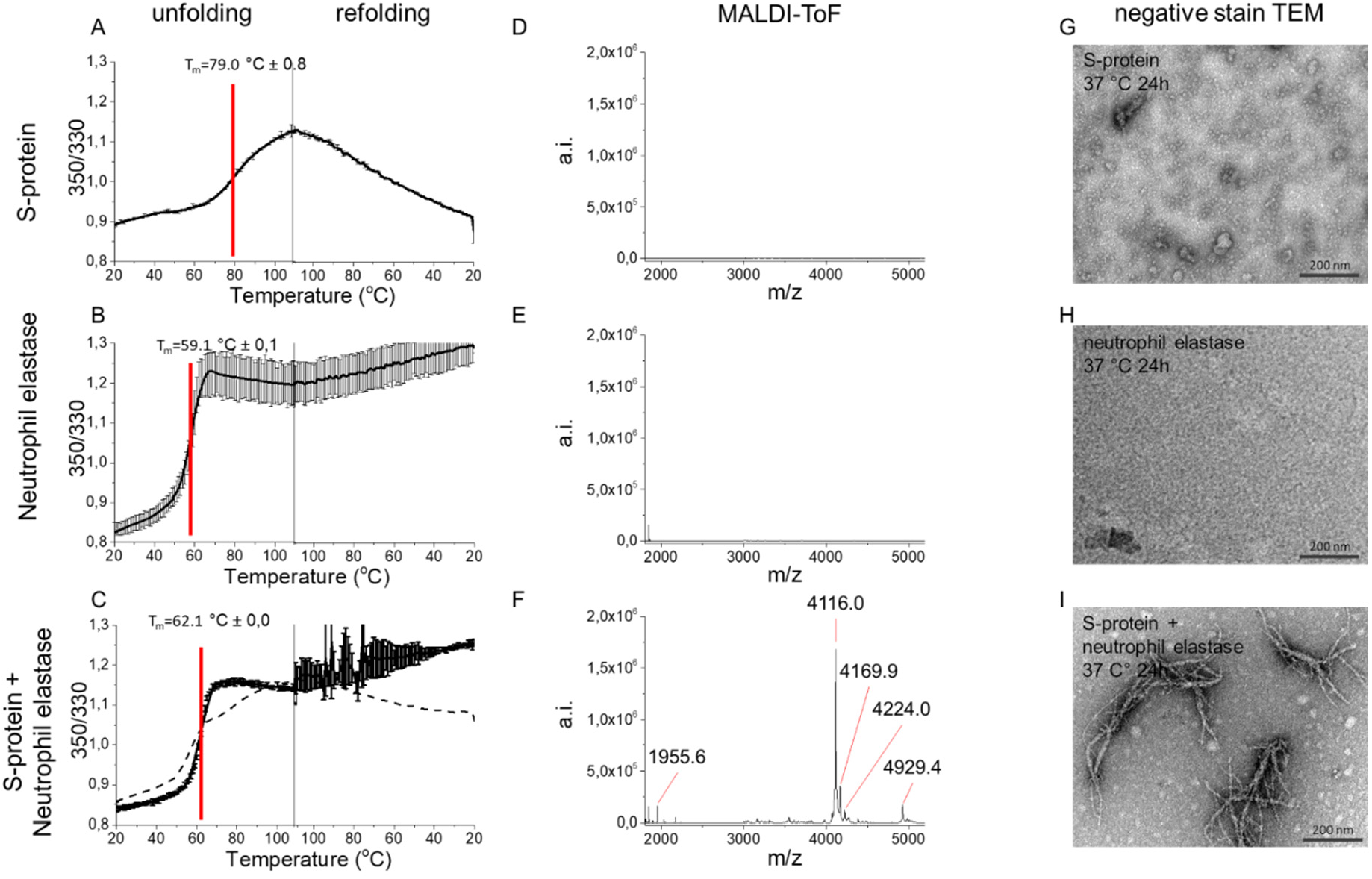
S-protein proteolysis by NE renders amyloid-like fibrils. A. Thermostability of the SARS-CoV-2 S-protein, B. NE and C. S-protein + NE, measured by DSF. The dashed line in C is the mathematical sum of S-protein and NE respectively from A and B which does not fit the experimental data, supporting cleavage of S-protein by NE. MALDI-ToF spectra of C18 isolated peptides of D. S-protein, E. NE, and F. S-protein + NE (6 h, 37 °C). TEM micrographs of G. S-protein alone depicting the expected trimers [17] H. NE alone, and I. S-protein and NE co-incubated at pH 8.4 for 24 h, 37 °C. Clusters of amyloidlike branched fibrils were formed in the co-incubation experiment in I.

S-protein showed a complex unfolding trajectory with multiple transitions around 45-65 °C, and a major unfolding transition with a midpoint of thermal denaturation (T_m_) 79 °C (Fig. 4A, SFig 3A), similar to that reported for another full-length S-protein construct [17]. S-protein refolded upon lowering the temperature albeit with a non-cooperative refolding transition (Fig. 4A). NE unfolded irreversibly with at T_m_ of 59 °C (Fig. 4B). The incubated S-protein + NE only showed an obvious transition for NE and did not refold upon lowering the temperature suggesting that S-protein had been cleaved by NE (Fig. 4C). We verified digestion by mass spectrometry where only the S-protein + NE experiment revealed peptide peaks (Fig. 4D-F). Most importantly we discovered amyloid-like fibril formation dependent on this proteolytic cleavage by TEM. Neither NE nor SARS-CoV-2 S-protein incubated alone formed fibrils (Fig. 4G-H). Fibrils were only found after co-incubation of the two proteins (Fig. 4I). The fibrils had an unusual morphology with evident branching (Fig. 4I). This morphology suggests involvement of proteolytically nicked S-protein within the fibril rendering nodes for branching of different amyloidogenic sequences (Fig. 4I, SFig. 4). To understand how NE cleaved S-protein we performed LC-MS/MS analysis of peptides formed after 1 min and 6 h of digestion at 2:1 excess of NE over S-protein. We mapped 98 identified NE cleavage peptides (STable 1) on the S-protein structure and classed these into three groups: i) formed after 1 min (Fig. 5A, B), ii) formed after 1 min and still present after 6 h (Fig. 5A, C), and iii) only present after 6 h (Fig. 5A, D). Initial cleavage and further digestion (group i) occurred mainly within the S2 domain with abundant cleavage of the HR domains and in the C-terminal part of S1. NTD and RBD were much less affected (Fig. 5A). Three persistent peptides formed after initial cleavage (group ii) originated from NTD and RBD. Several peptides only formed after 6 h of incubation (group iii). Strikingly the peptide from the segment 193-202 (FKNIDGYFKI, included in Spike191) was part of this group and was highly abundant after 6 h (STable 2). Compared to the amyloidogenic sequences three peptides containing segments from these were formed as free peptides (Spike191, Spike259, Spike1165) still present after 6 hours of co-incubation, two were digested early and disappeared (Spike532, Spike689), and two were likely still resident in the parent nicked S-protein (Spike362, Spike599). Hence the observed formed branched fibrils is likely composed of a mix of fibrils initiated by an amyloidogenic peptide seed recruiting nicked S-protein for elongation and branching.

**Figure 5.**
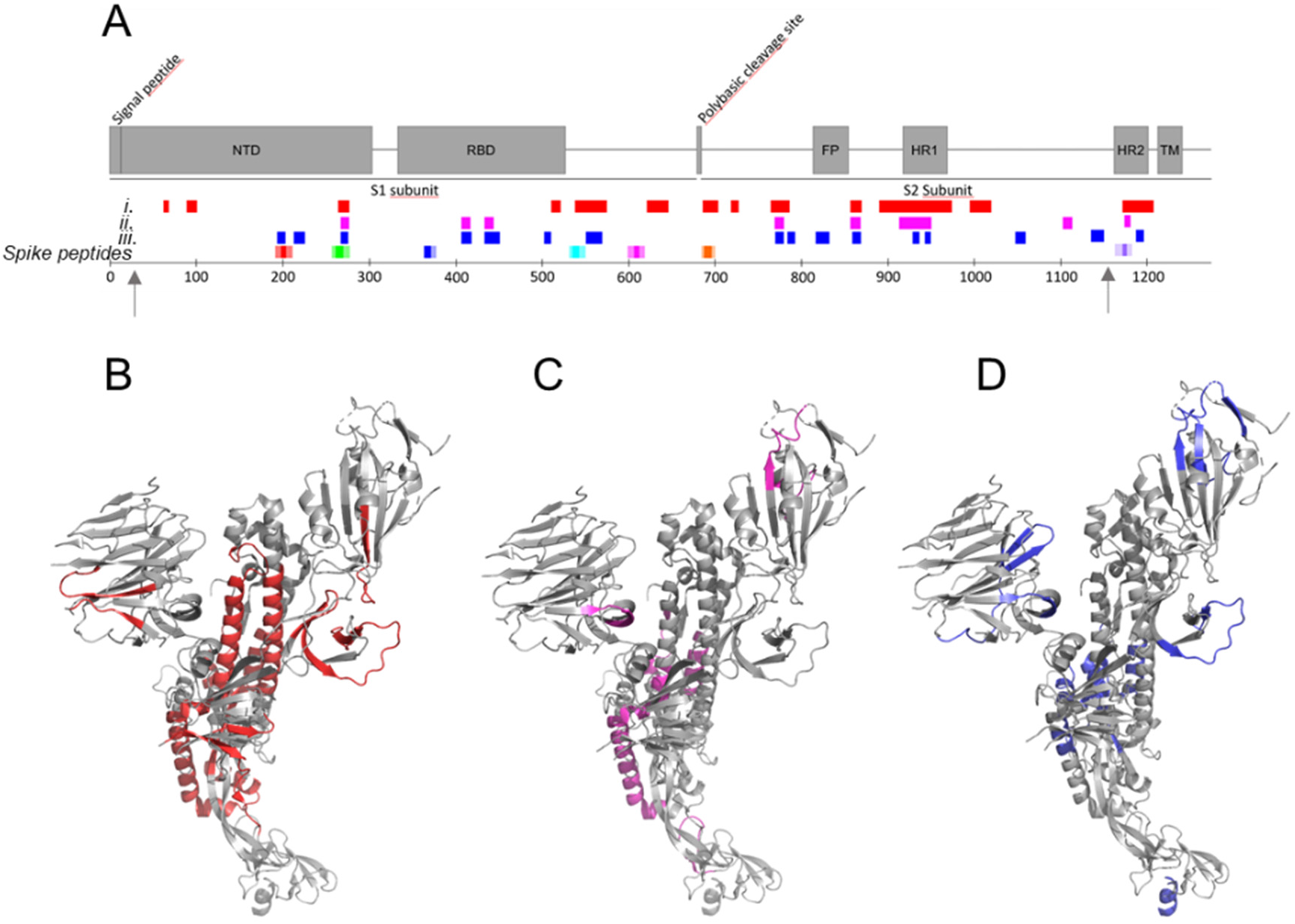
NE cleavage sites within full-length folded S-protein. A. Overall map of peptides identified by LC-MS/MS in correlation to the amyloidogenic Spike peptides (c.f. Fig. 1 and Table 1) and the S-protein domain structure. Arrows indicate the limits of the cryo-EM structure 27-1146. After i) 1 min (red), ii) 1 min and still persistent at 6 h (magenta), iii) only present after 6 h of incubation (blue). B-D. the same color code for cleaved peptide group i-iii mapped onto the protomer Cryo-EM S-protein structure PDB code 6VXX.

What are the implications of our findings? COVID-19 pathogenesis is multifactorial and complex [18]. Severe COVID-19 include acute respiratory distress syndrome (ARDS) from severe innate immune system inflammatory reactions resulting in lung damage [19]; Cytokine storm [20]; Heart damage, including inflammation of the heart muscle; Kidney damage; Neurological damage; Damage to the circulatory system resulting in poor blood flow; Long-COVID symptoms include persistent emotional illness and other mental health conditions resembling neurodegenerative diseases [18].

It has been proposed that severe inflammatory disease including ARDS in combination with SARS-CoV-2 protein aggregation might induce systemic AA amyloidosis [21]. Neurotropic colonization and cross-seeding of S-protein amyloid fibrils to induce aggregation of human endogenous proteins has been discussed in the context of neurodegeneration [4]. Notably, blood clotting associated with extra-cellular amyloidotic fibrillar aggregates in the blood stream have been reported in affected COVID-19 patients [22]. Hypercoagulation and impaired fibrinolysis were demonstrated in experimentally S-protein spiked blood plasma from healthy donors [22]. Similarly, amyloidosis is associated with cerebral amyloid angiopathy, blood coagulation disruption, fibrinolytic disturbance [23, 24] and FXII Kallikrein/Kinin activation and thromboinflammation [25], suggesting potential links between amyloidogenesis of S-protein and COVID-19 phenotypes.

In conclusion, we herein proposed a rather simple molecular mechanism for how SARS-CoV-2 S-protein endoproteolysis by NE can render exceptionally amyloidogenic S-peptides such as segment 193-202, and exposure of multiple amyloidogenic segments in proteolytically nicked S-protein. It is possible that other amyloidogenic peptides and S-protein nicked by other proteases could be involved. We found that all common coronaviruses infecting humans contain amyloidogenic sequences (SFig. 5A). Nonetheless, the magnitude of the diverse COVID-19 symptoms was not previously reported. The segment 193-212 is unique for and SARS-CoV-2 (SFig. 5B), which in combination with acute inflammation could explain the putative COVID-19 associated amyloidosis.

Furthermore, over the course of the year 2021 over 8 billion doses of COVID-19 vaccines have been administered worldwide. Most doses have provided SARS-CoV-2 S-protein as main antigen. A recent case report describes the serendipitous discovery of amyloid formation in a human patient locally at the site of vaccination and in a proximal lymph node within 24 hours of the first dose of mRNA vaccine coding for S-protein [26]. Neutrophil recruitment and activation at the site of vaccination is expected. The localized amyloid was detected by the Aβ amyloid PET tracer ^18^F-Florbetaben also known to bind to AL, AA and ATTR cardiac amyloid fibrils. The reactant alleged amyloidogenic protein in the vaccinated patient was not identified. We tested the fluorescent amyloid ligands CN-PiB (fluorescent benzothiazole analogue of Pittsburgh compound B) and DF-9 (fluorescent stilbene analogue of Florbetaben) and found strong fluorescence response towards Spike191 fibrils *in vitro* (SFig. 6), supporting the possibility that vaccine induced S-protein derived amyloid deposition was detected in the human PET imaging case study [26].

If the herein proposed mechanism for S-protein amyloid formation is associated with reported cardiac,- blood,- and nervous system disorders in certain vaccinated individuals [27] is not known, neither are the long-term consequences, but is strongly recommended to be investigated in this context.

## Supporting information

Supplemental Info Figs Tables Nystrom Hammarstrom

## AUTHOR INFORMATION

### Author Contributions

‡The research was performed, and manuscript was written with equal contributions of both authors.

Both authors have given approval to the final version of the manuscript.

## ACKNOWLEDGMENT

We thank Björn Wallner for generating Alpha fold 2 folding predictions of the synthetic peptides, Xiongyu Wu for synthesis of CN-PiB, and Jun Zhang for synthesis of DF-9. We acknowledge the use LiU core-facilities ProLinC, Medical Faculty Microscopy and Proteomics core especially Maria Turkina for collection of experimental data.

## ABBREVIATIONS

CNS: central nervous system
SARS-CoV-2: Severe Acute Respiratory Syndrome Coronavirus-2
S-protein: Spike protein
ThT,: Thioflavin T
CN-PiB: Cyano-Pittsburgh compound B
TEM: transmission electron microscopy
DSF: Differential scanning fluorimetry

