## Supplemental Info Figs Tables Nystrom Hammarstrom for "Amyloidogenesis of SARS-CoV-2 Spike Protein"

Nyström S, Hammarström P

**Supporting information 1: Experimental procedures**

**Solubilization and fibrillation of bulk peptide library**

A bulk library of 15-mer peptides with 11 amino acid overlap was purchased from GenScript, Netherlands (cat no catalog no RP30020), (Supporting information 2). The peptides were delivered as lyophilized powder in two subpools. Each subpool was treated separately as follows.

The lyophilized powder was reconstituted in autoclaved, ice-cold PBS (0.14 M NaCl, 0.0027 M KCl, 0.01 M $PO_4^{3-}$ (Medicago)) to a total concentration of 2 mg/ml peptide mixture (~12 µg/ml per peptide) and subsequently sonicated in water bath for 15 minutes. The resulting turbid mixture was transferred to centrifugation microtube and spun at 20 000 g at 4 °C for 20 minutes. The supernatant (hereafter called soluble fraction) was transferred to a fresh tube and the pellet was supplemented with hexafluoroisopropanol (HFIP) at a volume corresponding to 10 % of the initially added PBS (Hereafter called insoluble fraction). This resulted in a clear solution.

The soluble fraction was supplemented with 10% HFIP to match HFI concentration on the insoluble fraction. PBS was added to the insoluble fraction to reach a final concentration of 10% HFIP. 400 µl of each of the four preparations were aliquoted to tightly sealed glass vials and placed at 37 °C for 24 hours.

**Selection of amyloid prone segments**

Custom made peptides were selected from the full-length SARS-CoV-2 spike protein (protein ID: P0DTC2) with the aim to generate amyloid prone peptides. The primary sequence of the protein was subjected to the WALTZ algorithm (https://waltz.switchlab.org/ ) [1]. The algorithm was set to High selectivity to avoid false positives. The algorithm predicted 7 amyloidogenic sequences comprising 5-9 amino acids respectively (supporting information 3). The WALTZ predicted sequence was placed in the middle and flanking amino acids were selected to include hydrophobic and exclude charged residues. The peptides were named based on the number of the first amino acid in the respective sequence. Based on this information we selected peptides stretching over 20 amino acids. Due to solubility issues for some peptides, charged residues were reincluded and in some instances (Spike362 and Spike689) additional lysines and glycines were included to generate producible peptides. Spike259 was produced with 21 amino acids to allow for both the N-terminal tryptophan and the C-terminal lysine to be included.

**Solubilization and fibrillation of custom-made peptides**

Custom-made peptides (main text Table 1) were delivered from GenScript, Netherlands as lyophilized powder. 100% HFIP was added to each peptide to reach a final peptide concentration of 10 mg/ml. The vials were sonicated in water bath for 15 minutes and subsequently frozen at -20 °C until needed.

On day of fibrillation, the solubilized peptides were thawed at room temperature and diluted in HFIP to reach a peptide concentration of 1 mg/ml and sonicated in water bath 15 minutes. Triplicate samples for each peptide were prepared by adding HFIP solubilized peptide to ice cold phosphate buffered saline (PBS) pH 7.4, supplemented with ThT to reach a final concentration of 0,1 mg/ml peptide, 2 µM ThT and 10% HFIP. MALDI-ToF mass spectra were collected on the dissolved peptides to verify purity (SFig. 2.) Peptides at a concentration of 0.1 mg/ml dissolved in PBS was mixed 50/50 with alpha-Cyano-4-hydroxycinnamic acid matrix and were analyzed by UltrafleXtreme MALDI system (Bruker Daltonics, Bremen, Germany). A mixture of all peptides was also prepared by mixing equal volumes of each peptide preparation.

This renders a total peptide concentration of 0.1 mg/ml and a concentration of 0.014 mg/ml of each constituting peptide. The samples were distributed in 96-well black untreated half area plates with transparent bottom (Corning costar 3880) placed on a bed of ice. The plate was firmly sealed with aluminum and plastic sealing film to prevent evaporation of HFIP. The plate was placed at 37 °C in a Tecan Infinity Pro1000 plate reader with linear shaking between measurements with amplitude 2 mm and frequency 654 rpm. ThT intensity was monitored by bottom read mode with excitation at 440 nm and emission at 500 nm every 5 minutes for 24 hours (Main text Fig. 2).

Corresponding samples but with ThT omitted was also included on the plate for upstream experiments (below referred to as unstained reactions).

**Transmission electron microscopy**

TEM grids from each of the samples were prepared from the unstained fibrillation reactions as follows: Five µL of samples were placed on carbon coated copper grids (Carbon-B, Ted Pella Inc.) and were incubated for 2 minutes. Excessive salt was removed with one wash of 5 µL deionized water and grids were negatively stained with 2% uranyl acetate for 30 s. Grids were blotted dry and air dried overnight.

Transmission electron microscopy (TEM) imaging was performed using a Jeol JEM1400 Flash TEM microscope operating at 80 kV (Main text Fig. 2, 3, 4, SFig. 1 and SFig. 4).

**Congo red birefringence**

Aliquots from the unstained fibrillation reactions were prepared for evaluation of Congo red birefringence as follows: Five µl of Congo red stock solution (100 µM in milliQ water) was added to 45 µl of 0.1 mg/ml Spike-peptide fibrils in solution resulting in a molar ratio of Spike-peptide:dye 4:1. Stained fibrils were left to self-sediment over-night at 20 °C. 5 µl from the bottom of the pelleted samples were transferred to superfrost gold glass slides (Thermo Fisher, Walldorf, Germany) and allowed to dry. The dried droplets were covered with fluorescence mounting medium (Dako, Glosrup, Denmark). Congo red stained samples were analyzed using a Nikon light microscope equipped with polarizers for both incoming light and in front of the detector.

**Elastase digestion of full-length S-protein**

Full-length wild-type SARS-CoV-2 S-protein expressed in human HEK293 cells (Sigma-Aldrich catalogue # AGX819) was subjected to proteolytic cleavage by Human elastase expressed in human leucocytes (Sigma-Aldrich, catalogue # 324681) as follows:

The S-protein was delivered in Tris buffer pH 8 at a concentration of 5.8 µM (on a monomer basis). Elastase was delivered as lyophilized powder and was reconstituted in 10 mM Tris-HCl pH 8.4 (Tris-buffer) to a final concentration of 13.5 µM. Three sample types were prepared; 1 Spike diluted 1:1 in Tris-buffer; 2 Elastase diluted 1:1 in Tris buffer; 3 Spike and Elastase stocks mixed 1:1. All samples were incubated at 37 °C for 24 hours. After 24 hours PMSF was added to all samples to a final concentration of 1 mM. TEM grids were prepared, and TEM analysis was performed as described above.

**Thermal stability of elastase digested S-protein**

Thermal unfolding and refolding of SARS-CoV-2 S-protein in the presence or absence of elastase was assayed to verify folded protein from initiation and lack of refolding if nicking by protelolysis of S-protein was occurring. Samples were prepared with recombinant full-length SARS-CoV-2 S-protein 1 µM on a monomer basis (S-protein contains 12 Trp per monomer) in the presence or absence of neutrophil elastase 2.5 µM (3 Trp). Elastase alone was also assayed for comparison. All samples were prepared in 10 mM Tris-HCl buffer pH 8.4. Thermal stability scan was initiated 15 min after mixing of samples at 20 °C. Analysis was performed by Trp fluorescence by nanoscale differential fluorimetry (350nm/330 nm emission ratio) using the Prometehus instrumentation (Nanotemper). Settings: Thermal scan, 1 °C/min, 20-110 °C för unfolding and followed by refolding at 1 °C/min (Main text Fig. 4, SFig 3).

**Mass spectrometry of elastase digested S-protein**
Samples from the same reactions as for DSF were incubated in parallel at 37 °C for 6 h prior to preparation of samples for mass spectrometry. The reaction was stopped by mixing with equal volumes of 6 M GuHCl, 0.2 % TFA to a final concentration of 3 M GuHCl and 0.1% TFA. Samples were Zip-tipped using C4 and C18 Zip-tips respectively as described by the manufacturer (Millipore). Peptides isolated by C18 were mixed 50/50 with alpha-Cyano-4-hydroxycinnamic acid matrix and protein samples zip tipped with C4 were mixed with sinapinic acid as matrix for MALDI-ToF. Protein mass spectra were acquired on an UltrafleXtreme MALDI-TOF mass spectrometer (Bruker Daltonics Bremen Germany) instrument operated in the linear positive ion mode with flexControl software (Version 3.4, Bruker Daltonics). The MS spectra obtained were analyzed using flexAnalysis software (Version 3.4, Bruker Daltonics). (Main text Fig. 4, SFig. 2). No peaks were recovered above 50000 Da from the acquired spectra even for C4 isolated proteins. Elastase, were added, showed a peak with a centroid at 27100 ± 3 Da.

**LC–MS/MS and peptide identification**
The S-protein at a final concentration of 1.45 µM (on a monomer basis) was co-incubated with elastase at a final concentration of 2.7 µM in 10 mM Tris-HCl pH 8.4 (Tris-buffer) for 1 min or 6 h at 37 °C. Undigested S-protein was run as a control to check for possible *in vitro* degradation peptides. The reaction was stopped by mixing with equal volumes of 6 M GuHCl, 0.2 % TFA to a final concentration of 3 M GuHCl and 0.1% TFA. Samples were desalted using Ziptips C18 as described by the manufacturer (Millipore). Samples were eluted using 50% and 70% acetonitrile with 0.1 % TFA and vacuum dried using Speedvac concentrator (Thermo Scientific Savant).
The samples were redissolved in 0.1% formic acid in milli-Q water and were applied to an EASY-nanoLC II system (Thermo Fisher Scientific) with a C18 reverse chromatography column 20 mm × 100 µm C18 pre column followed by a 100 mm × 75 µm C18 column with particle size 5 µm (NanoSeparations, Nieuwkoop, Netherlands) and peptides were saperated at a flow rate of 300 nL/min by a gradient of 0.1% formic acid in water (A) and 0.1% formic acid in acetonitrile (B) as follows: from 2% B to 30% B in 60 minutes; from 30% B to 100% B in 60 minutes.

Automated online analyses were performed in positive mode by LTQ Orbitrap Velos Pro hybrid mass spectrometer (Thermo Scientific) equipped with a nano-electrospray source with Xcalibur software (v.2.6, Thermo Scientific). Full MS scans were collected with a range of 350–1800 *m*/*z*, a resolution of 30 000 (*m*/*z* 200), the top 20 most intense multiple charged ions were selected with an isolation window of 2.0 and fragmented in the linear ion trap by collision-induced dissociation with normalized collision energy of 35%. Dynamic exclusion was enabled ensuring peaks selected for fragmentation were excluded for 60 s.
Peptides were identified using using Sequest HT in Proteome Discoverer (Thermo Fisher Scientific, San Jose, CS version 2.5.0.400) and the cRAP database (cRAP protein sequences (thegpm.org)) merged with - P0DTC2 sequence for the SARS-CoV-2 S-protein (UniProtKB). and elastase cleavage prediction for peptide identification allowing missed cleavage sites. The following search parameters were used: elastase as a digestion enzyme; maximum number of missed cleavages 8; fragment ion mass tolerance 0.10 Da; parent ion mass tolerance 15.0 ppm.

SARS-CoV-2 S-protein peptides were quantified using two methods: peak intensity and spectral counts. Based on this, abundances of peptides originating from SARS-CoV-2 S-protein were compared for the 3 S-protein samples: undigested, incubated for 1 min with elastase, incubated for 6 h with elastase (Supporting Table 1_1 min vs 6 h Elastase_peptides and supporting Table 2_Quantification of abundant peaks).

##### Staining with fluorescent PET amyloid ligand analogues CN-PiB and DF-9

Unstained fibrils of Spike191 formed at 1 mg/ml was dilited to 0.1 mg/ml in PBS buffer pH 7.4 and were stained with a final concentration of 1 µM CN-PiB or DF-9 over night. 3 µl from the bottom of the sedimented samples were transferred to superfrost gold glass slides (Thermo Fisher, Walldorf, Germany) and allowed to dry. The dried droplets were covered with fluorescence mounting medium (Dako, Glosrup, Denmark). Stained samples were analyzed using a Leica600DM epifluorescence microscope equipped with long band pass filters and a hyperspectral camera (SpectraCube, ASI, Israel). Settings: excitation 350 nm emission 400-700 nm, 25 ms exposure, 20x objective. (SFig. 6).

##### Supporting information 2: Bulk S-protein peptide library sequences

Marked in yellow are peptides marked as amyloidogenic based on WALTZ "best overall performance" settings [1].

Red letters were indicated as amyloidogenic with WALTZ "high specificity" settings.

**Subpool 1**

| | |
|---|---|
| SARS-CoV-2 Spike_1 | MFVFLVLLPLVSSQC |
| SARS-CoV-2 Spike_2 | LVLLPLVSSQCVNLT |
| SARS-CoV-2 Spike_3 | PLVSSQCVNLTTRTQ |
| SARS-CoV-2 Spike_4 | SQCVNLTTRTQLPPA |
| SARS-CoV-2 Spike_5 | NLTTRTQLPPAYTNS |
| SARS-CoV-2 Spike_6 | RTQLPPAYTNSFTRG |
| SARS-CoV-2 Spike_7 | PPAYTNSFTRGVYYP |
| SARS-CoV-2 Spike_8 | TNSFTRGVYYPDKVF |
| SARS-CoV-2 Spike_9 | TRGVYYPDKVFRSSV |
| SARS-CoV-2 Spike_10 | YYPDKVFRSSVLHST |
| SARS-CoV-2 Spike_11 | KVFRSSVLHSTQDLF |
| SARS-CoV-2 Spike_12 | SSVLHSTQDLFLPFF |
| SARS-CoV-2 Spike_13 | HSTQDLFLPFFSNVT |
| SARS-CoV-2 Spike_14 | DLFLPFFSNVTWFHA |
| SARS-CoV-2 Spike_15 | PFFSNVTWFHAIHVS |
| SARS-CoV-2 Spike_16 | NVTWFHAIHVSGTNG |
| SARS-CoV-2 Spike_17 | FHAIHVSGTNGTKRF |
| SARS-CoV-2 Spike_18 | HVSGTNGTKRFDNPV |
| SARS-CoV-2 Spike_19 | TNGTKRFDNPVLPFN |
| SARS-CoV-2 Spike_20 | KRFDNPVLPFNDGVY |
| SARS-CoV-2 Spike_21 | NPVLPFNDGVYFAST |
| SARS-CoV-2 Spike_22 | PFNDGVYFASTEKSN |
| SARS-CoV-2 Spike_23 | GVYFASTEKSNIIRG |
| SARS-CoV-2 Spike_24 | ASTEKSNIIRGWIFG |
| SARS-CoV-2 Spike_25 | KSNIIRGWIFGTTLD |
| SARS-CoV-2 Spike_26 | IRGWIFGTTLDSKTQ |
| SARS-CoV-2 Spike_27 | IFGTTLDSKTQSLLI |
| SARS-CoV-2 Spike_28 | TLDSKTQSLLIVNNA |
| SARS-CoV-2 Spike_29 | KTQSLLIVNNATNVV |
| SARS-CoV-2 Spike_30 | LLIVNNATNVVIKVC |
| SARS-CoV-2 Spike_31 | NNATNVVIKVCEFQF |
| SARS-CoV-2 Spike_32 | NVVIKVCEFQFCNDP |
| SARS-CoV-2 Spike_33 | KVCEFQFCNDPFLGV |
| SARS-CoV-2 Spike_34 | FQFCNDPFLGVYYHK |
| SARS-CoV-2 Spike_35 | NDPFLGVYYHKNNKS |
| SARS-CoV-2 Spike_36 | LGVYYHKNNKSWMES |
| SARS-CoV-2 Spike_37 | YHKNNKSWMESEFRV |
| SARS-CoV-2 Spike_38 | NKSWMESEFRVYSSA |
| SARS-CoV-2 Spike_39 | MESEFRVYSSANNCT |
| SARS-CoV-2 Spike_40 | FRVYSSANNCTFEYV |
| SARS-CoV-2 Spike_41 | SSANNCTFEYVSQPF |
| SARS-CoV-2 Spike_42 | NCTFEYVSQPFLMDL |
| SARS-CoV-2 Spike_43 | EYVSQPFLMDLEGKQ |
| SARS-CoV-2 Spike_44 | QPFLMDLEGKQGNFK |
| SARS-CoV-2 Spike_45 | MDLEGKQGNFKNLRE |
| SARS-CoV-2 Spike_46 | GKQGNFKNLREFVFK |
| SARS-CoV-2 Spike_47 | NFKNLREFVFKNIDG |
| SARS-CoV-2 Spike_48 | LREFVFKNIDGYFKI |
| SARS-CoV-2 Spike_49 | VFKNIDGYFKIYSKH |
| SARS-CoV-2 Spike_50 | IDGYFKIYSKHTPIN |
| SARS-CoV-2 Spike_51 | FKIYSKHTPINLVRD |
| SARS-CoV-2 Spike_52 | SKHTPINLVRDLPQG |
| SARS-CoV-2 Spike_53 | PINLVRDLPQGFSAL |
| SARS-CoV-2 Spike_54 | VRDLPQGFSALEPLV |
| SARS-CoV-2 Spike_55 | PQGFSALEPLVDLPI |
| SARS-CoV-2 Spike_56 | SALEPLVDLPIGINI |
| SARS-CoV-2 Spike_57 | PLVDLPIGINITRFQ |
| SARS-CoV-2 Spike_58 | LPIGINITRFQTLLA |
| SARS-CoV-2 Spike_59 | INITRFQTLLALHRS |
| SARS-CoV-2 Spike_60 | RFQTLLALHRSYLTP |
| SARS-CoV-2 Spike_61 | LLALHRSYLTPGDSS |
| SARS-CoV-2 Spike_62 | HRSYLTPGDSSSGWT |
| SARS-CoV-2 Spike_63 | LTPGDSSSGWTAGAA |
| SARS-CoV-2 Spike_64 | DSSSGWTAGAAAYYV |
| SARS-CoV-2 Spike_65 | GWTAGAAAYYVGYLQ |
| SARS-CoV-2 Spike_66 | GAAAYYVGYLQPRTF |
| SARS-CoV-2 Spike_67 | YYVGYLQPRTFLLKY |
| SARS-CoV-2 Spike_68 | YLQPRTFLLKYNENG |
| SARS-CoV-2 Spike_69 | RTFLLKYNENGTITD |
| SARS-CoV-2 Spike_70 | LKYNENGTITDAVDC |
| SARS-CoV-2 Spike_71 | ENGTITDAVDCALDP |
| SARS-CoV-2 Spike_72 | ITDAVDCALDPLSET |
| SARS-CoV-2 Spike_73 | VDCALDPLSETKCTL |
| SARS-CoV-2 Spike_74 | LDPLSETKCTLKSFT |
| SARS-CoV-2 Spike_75 | SETKCTLKSFTVEKG |
| SARS-CoV-2 Spike_76 | CTLKSFTVEKGIYQT |
| SARS-CoV-2 Spike_77 | SFTVEKGIYQTSNFR |
| SARS-CoV-2 Spike_78 | EKGIYQTSNFRVQPT |
| SARS-CoV-2 Spike_79 | YQTSNFRVQPTESIV |
| SARS-CoV-2 Spike_80 | NFRVQPTESIVRFPN |
| SARS-CoV-2 Spike_81 | QPTESIVRFPNITNL |
| SARS-CoV-2 Spike_82 | SIVRFPNITNLCPFG |
| SARS-CoV-2 Spike_83 | FPNITNLCPFGEVFN |
| SARS-CoV-2 Spike_84 | TNLCPFGEVFNATRF |
| SARS-CoV-2 Spike_85 | PFGEVFNATRFASVY |
| SARS-CoV-2 Spike_86 | VFNATRFASVYAWNR |
| SARS-CoV-2 Spike_87 | TRFASVYAWNRKRIS |
| SARS-CoV-2 Spike_88 | SVYAWNRKRISNCVA |
| SARS-CoV-2 Spike_89 | WNRKRISNCVADYSV |
| SARS-CoV-2 Spike_90 | RISNCVADYSVLYNS |
| SARS-CoV-2 Spike_91 | CVADYSVLYNSASFS |
| SARS-CoV-2 Spike_92 | YSVLYNSASFSTFKC |

| | |
|---|---|
| SARS-CoV-2 Spike_93 | YNSASFSTFKCYGVS |
| SARS-CoV-2 Spike_94 | SFSTFKCYGVSPTKL |
| SARS-CoV-2 Spike_95 | FKCYGVSPTKLNDLC |
| SARS-CoV-2 Spike_96 | GVSPTKLNDLCFTNV |
| SARS-CoV-2 Spike_97 | TKLNDLCFTNVYADS |
| SARS-CoV-2 Spike_98 | DLCFTNVYADSFVIR |
| SARS-CoV-2 Spike_99 | TNVYADSFVIRGDEV |
| SARS-CoV-2 Spike_100 | ADSFVIRGDEVRQIA |
| SARS-CoV-2 Spike_101 | VIRGDEVRQIAPGQT |
| SARS-CoV-2 Spike_102 | DEVRQIAPGQTGKIA |
| SARS-CoV-2 Spike_103 | QIAPGQTGKIADYNY |
| SARS-CoV-2 Spike_104 | GQTGKIADYNYKLPD |
| SARS-CoV-2 Spike_105 | KIADYNYKLPDDFTG |
| SARS-CoV-2 Spike_106 | YNYKLPDDFTGCVIA |
| SARS-CoV-2 Spike_107 | LPDDFTGCVIAWNSN |
| SARS-CoV-2 Spike_108 | FTGCVIAWNSNNLDS |
| SARS-CoV-2 Spike_109 | VIAWNSNNLDSKVGG |
| SARS-CoV-2 Spike_110 | NSNNLDSKVGGNYNY |
| SARS-CoV-2 Spike_111 | LDSKVGGNYNYLYRL |
| SARS-CoV-2 Spike_112 | VGGNYNYLYRLFRKS |
| SARS-CoV-2 Spike_113 | YNYLYRLFRKSNLKP |
| SARS-CoV-2 Spike_114 | YRLFRKSNLKPFERD |
| SARS-CoV-2 Spike_115 | RKSNLKPFERDISTE |
| SARS-CoV-2 Spike_116 | LKPFERDISTEIYQA |
| SARS-CoV-2 Spike_117 | ERDISTEIYQAGSTP |
| SARS-CoV-2 Spike_118 | STEIYQAGSTPCNGV |
| SARS-CoV-2 Spike_119 | YQAGSTPCNGVEGFN |
| SARS-CoV-2 Spike_120 | STPCNGVEGFNCYFP |
| SARS-CoV-2 Spike_121 | NGVEGFNCYFPLQSY |
| SARS-CoV-2 Spike_122 | GFNCYFPLQSYGFQP |
| SARS-CoV-2 Spike_123 | YFPLQSYGFQPTNGV |
| SARS-CoV-2 Spike_124 | QSYGFQPTNGVGYQP |
| SARS-CoV-2 Spike_125 | FQPTNGVGYQPYRVV |
| SARS-CoV-2 Spike_126 | NGVGYQPYRVVVLSF |
| SARS-CoV-2 Spike_127 | YQPYRVVVLSFELLH |
| SARS-CoV-2 Spike_128 | RVVVLSFELLHAPAT |
| SARS-CoV-2 Spike_129 | LSFELLHAPATVCGP |
| SARS-CoV-2 Spike_130 | LLHAPATVCGPKKST |
| SARS-CoV-2 Spike_131 | PATVCGPKKSTNLVK |
| SARS-CoV-2 Spike_132 | CGPKKSTNLVKNKCV |
| SARS-CoV-2 Spike_133 | KSTNLVKNKCVNFNF |
| SARS-CoV-2 Spike_134 | LVKNKCVNFNFNGLT |
| SARS-CoV-2 Spike_135 | KCVNFNFNGLTGTGV |
| SARS-CoV-2 Spike_136 | FNFNGLTGTGVLTES |
| SARS-CoV-2 Spike_137 | GLTGTGVLTESNKKF |
| SARS-CoV-2 Spike_138 | TGVLTESNKKFLPFQ |
| SARS-CoV-2 Spike_139 | TESNKKFLPFQQFGR |
| SARS-CoV-2 Spike_140 | KKFLPFQQFGRDIAD |
| SARS-CoV-2 Spike_141 | PFQQFGRDIADTTDA |
| SARS-CoV-2 Spike_142 | FGRDIADTTDAVRDP |
| SARS-CoV-2 Spike_143 | IADTTDAVRDPQTLE |
| SARS-CoV-2 Spike_144 | TDAVRDPQTLEILDI |
| SARS-CoV-2 Spike_145 | RDPQTLEILDITPCS |
| SARS-CoV-2 Spike_146 | TLEILDITPCSFGGV |
| SARS-CoV-2 Spike_147 | LDITPCSFGGVSVIT |
| SARS-CoV-2 Spike_148 | PCSFGGVSVITPGTN |
| SARS-CoV-2 Spike_149 | GGVSVITPGTNTSNQ |
| SARS-CoV-2 Spike_150 | VITPGTNTSNQVAVL |
| SARS-CoV-2 Spike_151 | GTNTSNQVAVLYQDV |
| SARS-CoV-2 Spike_152 | SNQVAVLYQDVNCTE |
| SARS-CoV-2 Spike_153 | AVLYQDVNCTEVPVA |
| SARS-CoV-2 Spike_154 | QDVNCTEVPVAIHAD |
| SARS-CoV-2 Spike_155 | CTEVPVAIHADQLTP |
| SARS-CoV-2 Spike_156 | PVAIHADQLTPTWRV |
| SARS-CoV-2 Spike_157 | HADQLTPTWRVYSTG |
| SARS-CoV-2 Spike_158 | LTPTWRVYSTGSNVF |

**Subpool 2**

| | |
|---|---|
| SARS-CoV-2 Spike_159 | WRVYSTGSNVFQTRA |
| SARS-CoV-2 Spike_160 | STGSNVFQTRAGCLI |
| SARS-CoV-2 Spike_161 | NVFQTRAGCLIGAEH |
| SARS-CoV-2 Spike_162 | TRAGCLIGAEHVNNS |
| SARS-CoV-2 Spike_163 | CLIGAEHVNNSYECD |
| SARS-CoV-2 Spike_164 | AEHVNNSYECDIPIG |
| SARS-CoV-2 Spike_165 | NNSYECDIPIGAGIC |
| SARS-CoV-2 Spike_166 | ECDIPIGAGICASYQ |
| SARS-CoV-2 Spike_167 | PIGAGICASYQTQTN |
| SARS-CoV-2 Spike_168 | GICASYQTQTNSPRR |
| SARS-CoV-2 Spike_169 | SYQTQTNSPRRARSV |
| SARS-CoV-2 Spike_170 | QTNSPRRARSVASQS |
| SARS-CoV-2 Spike_171 | PRRARSVASQSIIAY |
| SARS-CoV-2 Spike_172 | RSVASQSIIAYTMSL |
| SARS-CoV-2 Spike_173 | SQSIIAYTMSLGAEN |
| SARS-CoV-2 Spike_174 | IAYTMSLGAENSVAY |
| SARS-CoV-2 Spike_175 | MSLGAENSVAYSNNS |
| SARS-CoV-2 Spike_176 | AENSVAYSNNSIAIP |
| SARS-CoV-2 Spike_177 | VAYSNNSIAIPTNFT |
| SARS-CoV-2 Spike_178 | NNSIAIPTNFTISVT |
| SARS-CoV-2 Spike_179 | AIPTNFTISVTTEIL |
| SARS-CoV-2 Spike_180 | NFTISVTTEILPVSM |
| SARS-CoV-2 Spike_181 | SVTTEILPVSMTKTS |
| SARS-CoV-2 Spike_182 | EILPVSMTKTSVDCT |
| SARS-CoV-2 Spike_183 | VSMTKTSVDCTMYIC |
| SARS-CoV-2 Spike_184 | KTSVDCTMYICGDST |
| SARS-CoV-2 Spike_185 | DCTMYICGDSTECSN |
| SARS-CoV-2 Spike_186 | YICGDSTECSNLLLQ |
| SARS-CoV-2 Spike_187 | DSTECSNLLLQYGSF |
| SARS-CoV-2 Spike_188 | CSNLLLQYGSFCTQL |
| SARS-CoV-2 Spike_189 | LLQYGSFCTQLNRAL |
| SARS-CoV-2 Spike_190 | GSFCTQLNRALTGIA |
| SARS-CoV-2 Spike_191 | TQLNRALTGIAVEQD |
| SARS-CoV-2 Spike_192 | RALTGIAVEQDKNTQ |
| SARS-CoV-2 Spike_193 | GIAVEQDKNTQEVFA |
| SARS-CoV-2 Spike_194 | EQDKNTQEVFAQVKQ |
| SARS-CoV-2 Spike_195 | NTQEVFAQVKQIYKT |
| SARS-CoV-2 Spike_196 | VFAQVKQIYKTPPIK |
| SARS-CoV-2 Spike_197 | VKQIYKTPPIKDFGG |
| SARS-CoV-2 Spike_198 | YKTPPIKDFGGFNFS |
| SARS-CoV-2 Spike_199 | PIKDFGGFNFSQILP |
| SARS-CoV-2 Spike_200 | FGGFNFSQILPDPSK |
| SARS-CoV-2 Spike_201 | NFSQILPDPSKPSKR |
| SARS-CoV-2 Spike_202 | ILPDPSKPSKRSFIE |
| SARS-CoV-2 Spike_203 | PSKPSKRSFIEDLLF |
| SARS-CoV-2 Spike_204 | SKRSFIEDLLFNKVT |
| SARS-CoV-2 Spike_205 | FIEDLLFNKVTLADA |
| SARS-CoV-2 Spike_206 | LLFNKVTLADAGFIK |
| SARS-CoV-2 Spike_207 | KVTLADAGFIKQYGD |
| SARS-CoV-2 Spike_208 | ADAGFIKQYGDCLGD |
| SARS-CoV-2 Spike_209 | FIKQYGDCLGDIAAR |
| SARS-CoV-2 Spike_210 | YGDCLGDIAARDLIC |
| SARS-CoV-2 Spike_211 | LGDIAARDLICAQKF |
| SARS-CoV-2 Spike_212 | AARDLICAQKFNGLT |
| SARS-CoV-2 Spike_213 | LICAQKFNGLTVLPP |
| SARS-CoV-2 Spike_214 | QKFNGLTVLPPLLTD |
| SARS-CoV-2 Spike_215 | GLTVLPPLLTDEMIA |
| SARS-CoV-2 Spike_216 | LPPLLTDEMIAQYTS |
| SARS-CoV-2 Spike_217 | LTDEMIAQYTSALLA |
| SARS-CoV-2 Spike_218 | MIAQYTSALLAGTIT |
| SARS-CoV-2 Spike_219 | YTSALLAGTITSGWT |
| SARS-CoV-2 Spike_220 | LLAGTITSGWTFGAG |
| SARS-CoV-2 Spike_221 | TITSGWTFGAGAALQ |
| SARS-CoV-2 Spike_222 | GWTFGAGAALQIPFA |
| SARS-CoV-2 Spike_223 | GAGAALQIPFAMQMA |
| SARS-CoV-2 Spike_224 | ALQIPFAMQMAYRFN |
| SARS-CoV-2 Spike_225 | PFAMQMAYRFNGIGV |
| SARS-CoV-2 Spike_226 | QMAYRFNGIGVTQNV |
| SARS-CoV-2 Spike_227 | RFNGIGVTQNVLYEN |
| SARS-CoV-2 Spike_228 | IGVTQNVLYENQKLI |
| SARS-CoV-2 Spike_229 | QNVLYENQKLIANQF |
| SARS-CoV-2 Spike_230 | YENQKLIANQFNSAI |
| SARS-CoV-2 Spike_231 | KLIANQFNSAIGKIQ |
| SARS-CoV-2 Spike_232 | NQFNSAIGKIQDSLS |
| SARS-CoV-2 Spike_233 | SAIGKIQDSLSSTAS |
| SARS-CoV-2 Spike_234 | KIQDSLSSTASALGK |
| SARS-CoV-2 Spike_235 | SLSSTASALGKLQDV |
| SARS-CoV-2 Spike_236 | TASALGKLQDVVNQN |
| SARS-CoV-2 Spike_237 | LGKLQDVVNQNAQAL |
| SARS-CoV-2 Spike_238 | QDVVNQNAQALNTLV |
| SARS-CoV-2 Spike_239 | NQNAQALNTLVKQLS |
| SARS-CoV-2 Spike_240 | QALNTLVKQLSSNFG |
| SARS-CoV-2 Spike_241 | TLVKQLSSNFGAISS |
| SARS-CoV-2 Spike_242 | QLSSNFGAISSVLND |
| SARS-CoV-2 Spike_243 | NFGAISSVLNDILSR |
| SARS-CoV-2 Spike_244 | ISSVLNDILSRLDKV |
| SARS-CoV-2 Spike_245 | LNDILSRLDKVEAEV |
| SARS-CoV-2 Spike_246 | LSRLDKVEAEVQIDR |
| SARS-CoV-2 Spike_247 | DKVEAEVQIDRLITG |
| SARS-CoV-2 Spike_248 | AEVQIDRLITGRLQS |
| SARS-CoV-2 Spike_249 | IDRLITGRLQSLQTY |
| SARS-CoV-2 Spike_250 | ITGRLQSLQTYVTQQ |
| SARS-CoV-2 Spike_251 | LQSLQTYVTQQLIRA |
| SARS-CoV-2 Spike_252 | QTYVTQQLIRAAEIR |
| SARS-CoV-2 Spike_253 | TQQLIRAAEIRASAN |
| SARS-CoV-2 Spike_254 | IRAAEIRASANLAAT |
| SARS-CoV-2 Spike_255 | EIRASANLAATKMSE |
| SARS-CoV-2 Spike_256 | SANLAATKMSECVLG |
| SARS-CoV-2 Spike_257 | AATKMSECVLGQSKR |
| SARS-CoV-2 Spike_258 | MSECVLGQSKRVDFC |
| SARS-CoV-2 Spike_259 | VLGQSKRVDFCGKGY |
| SARS-CoV-2 Spike_260 | SKRVDFCGKGYHLMS |
| SARS-CoV-2 Spike_261 | DFCGKGYHLMSFPQS |
| SARS-CoV-2 Spike_262 | KGYHLMSFPQSAPHG |
| SARS-CoV-2 Spike_263 | LMSFPQSAPHGVVFL |
| SARS-CoV-2 Spike_264 | PQSAPHGVVFLHVTY |
| SARS-CoV-2 Spike_265 | PHGVVFLHVTYVPAQ |

| | | |
|---|---|---|
| SARS-CoV-2 | Spike_266 | VFLHVTYVPAQEKNF |
| SARS-CoV-2 | Spike_267 | VTYVPAQEKNFTTAP |
| SARS-CoV-2 | Spike_268 | PAQEKNFTTAPAICH |
| SARS-CoV-2 | Spike_269 | KNFTTAPAICHDGKA |
| SARS-CoV-2 | Spike_270 | TAPAICHDGKAHFPR |
| SARS-CoV-2 | Spike_271 | ICHDGKAHFPREGVF |
| SARS-CoV-2 | Spike_272 | GKAHFPREGVFVSNG |
| SARS-CoV-2 | Spike_273 | FPREGVFVSNGTHWF |
| SARS-CoV-2 | Spike_274 | GVFVSNGTHWFVTQR |
| SARS-CoV-2 | Spike_275 | SNGTHWFVTQRNFYE |
| SARS-CoV-2 | Spike_276 | HWFVTQRNFYEPQII |
| SARS-CoV-2 | Spike_277 | TQRNFYEPQIITTDN |
| SARS-CoV-2 | Spike_278 | FYEPQIITTDNTFVS |
| SARS-CoV-2 | Spike_279 | QIITTDNTFVSGNCD |
| SARS-CoV-2 | Spike_280 | TDNTFVSGNCDVVIG |
| SARS-CoV-2 | Spike_281 | FVSGNCDVVIGIVNN |
| SARS-CoV-2 | Spike_282 | NCDVVIGIVNNTVYD |
| SARS-CoV-2 | Spike_283 | VIGIVNNTVYDPLQP |
| SARS-CoV-2 | Spike_284 | VNNTVYDPLQPELDS |
| SARS-CoV-2 | Spike_285 | VYDPLQPELDSFKEE |
| SARS-CoV-2 | Spike_286 | LQPELDSFKEELDKY |
| SARS-CoV-2 | Spike_287 | LDSFKEELDKYFKNH |
| SARS-CoV-2 | Spike_288 | KEELDKYFKNHTSPD |
| SARS-CoV-2 | Spike_289 | DKYFKNHTSPDVDLG |
| SARS-CoV-2 | Spike_290 | KNHTSPDVDLGDISG |
| SARS-CoV-2 | Spike_291 | SPDVDLGDISGINAS |
| SARS-CoV-2 | Spike_292 | DLGDISGINASVVNI |
| SARS-CoV-2 | Spike_293 | ISGINASVVNIQKEI |
| SARS-CoV-2 | Spike_294 | NASVVNIQKEIDRLN |
| SARS-CoV-2 | Spike_295 | VNIQKEIDRLNEVAK |
| SARS-CoV-2 | Spike_296 | KEIDRLNEVAKNLNE |
| SARS-CoV-2 | Spike_297 | RLNEVAKNLNESLID |
| SARS-CoV-2 | Spike_298 | VAKNLNESLIDLQEL |
| SARS-CoV-2 | Spike_299 | LNESLIDLQELGKYE |
| SARS-CoV-2 | Spike_300 | LIDLQELGKYEQYIK |
| SARS-CoV-2 | Spike_301 | QELGKYEQYIKWPWY |
| SARS-CoV-2 | Spike_302 | KYEQYIKWPWYIWLG |
| SARS-CoV-2 | Spike_303 | YIKWPWYIWLGFIAG |
| SARS-CoV-2 | Spike_304 | PWYIWLGFIAGLIAI |
| SARS-CoV-2 | Spike_305 | WLGFIAGLIAIVMVT |
| SARS-CoV-2 | Spike_306 | IAGLIAIVMVTIMLC |
| SARS-CoV-2 | Spike_307 | IAIVMVTIMLCCMTS |
| SARS-CoV-2 | Spike_308 | MVTIMLCCMTSCCSC |
| SARS-CoV-2 | Spike_309 | MLCCMTSCCSCLKGC |
| SARS-CoV-2 | Spike_310 | MTSCCSCLKGCCSCG |
| SARS-CoV-2 | Spike_311 | CSCLKGCCSCGSCCK |
| SARS-CoV-2 | Spike_312 | KGCCSCGSCCKFDED |
| SARS-CoV-2 | Spike_313 | SCGSCCKFDEDDSEP |
| SARS-CoV-2 | Spike_314 | CCKFDEDDSEPVLKG |
| SARS-CoV-2 | Spike_315 | DEDDSEPVLKGVKLH |
| SARS-CoV-2 | Spike_316 | SEPVLKGVKLHYT |

**Supporting information 3: Selection of amyloidogenic segments for peptide synthesis and comparison with peptides rendered by *in silico* elastase digestion**

WALTZ prediction "high specificity", pH 7.0 [1], on SARS-CoV-2 spike protein indicated with asterisk under the sequence. The synthesized peptide sequences highlighted in colors corresponding to main text Fig. 1. Segments highlighted in grey are peptides predicted to be the result of cleaving with Neutrophil elastase using Expasy peptide cutter and rendering peptides with 3 or more consecutive amino acids in one of the amyloidogenic regions.

>sp|P0DTC2|SPIKE_SARS2

MFVFLVLLPLVSSQCVNLTTRTQLPPAYTNSFTRGVYYPDKVFRSSVLHSTQDLFLPFFS

NVTWFHAIHVSGTNGTKRFDNPVLPFNDGVYFASTEKSNIIRGWIFGTTLDSKTQSLLIV

NNATNVVIKVCEFQFCNDPFLGVYYHKNNKSWMESEFRVYSSANNCTFEYVSQPFLMDLE

GKQGNFKNLRE**FVFKNIDGYFKIYSKHTPIN**LVRDLPQGFSALEPLVDLPIGINITRFQT

******

**FKNIDGYFKIYSKHTPINLV**

LLALHRSYLTPGDSSSG**WTAGAAAYYVGYLQPRTFLLK**YNENGTITDAVDCALDPLSETK

*********

**GYLQPRTFLLKYNENGTITDA**

CTLKSFTVEKGIYQTSNFRVQPTESIVRFPNITNLCPFGEVFNATRFASVYAWNRKRISN

CVAD**YSVLYNSASFSTFK**CYGVSPTKLNDLCFTNVYADSFVIRGDEVRQIAPGQTGKIAD

********

**LYNSA**

YNYKLPDDFTGCVIAWNSNNLDSKVGGNYNYLYRLFRKSNLKPFERDISTEIYQAGSTPC

NGVEGFNCYFPLQSYGFQPTNGVGYQPYRVVVLSFELLHAPATVCGPKKSTN**LVKNKCVN**

****

**N**

**FNFNGLTGTGV**LTESNKKFLPFQQFGRDIADTTDAVRDPQTLEILDITPCSFGGVSVITP

****

**FNFNGLTGTGV**

**GTNTSNQVAVLYQDVNCTEV**PVAIHADQLTPTWRVYSTGSNVFQTRAGCLIGAEHVNNSY

******

**LYQDV**

ECDIPIGAGICASYQTQTNSPRRAR**SVASQSIIAYTMSLGA**ENSVAYSNNSIAIPTNFTI

*********

**SQSIIA**

SVTTEILPVSMTKTSVDCTMYICGDSTECSNLLLQYGSFCTQLNRALTGIAVEQDKNTQE

VFAQVKQIYKTPPIKDFGGFNFSQILPDPSKPSKRSFIEDLLFNKVTLADAGFIKQYGDC

LGDIAARDLICAQKFNGLTVLPPLLTDEMIAQYTSALLAGTITSGWTFGAGAALQIPFAM

QMAYRFNGIGVTQNVLYENQKLIANQFNSAIGKIQDSLSSTASALGKLQDVVNQNAQALN

TLVKQLSSNFGAISSVLNDILSRLDKVEAEVQIDRLITGRLQSLQTYVTQQLIRAAEIRA

SANLAATKMSECVLGQSKRVDFCGKGYHLMSFPQSAPHGVVFLHVTYVPAQEKNFTTAPA

ICHDGKAHFPREGVFVSNGTHWFVTQRNFYEPQIITTDNTFVSGNCDVVIGIVNNTVYDP

LQPELDSFKEELDKYFKNHTSPDVD**LGDISGINASVVNIQKEIDR**LNEVAKNLNESLIDL

*****

QELGKYEQYIKWPWYIWLGFIAGLIAIVMVTIMLCCMTSCCSCLKGCCSCGSCCKFDEDD

SEPVLKGVKLHYT

**Supporting Figure 1: Fibrils formed from bulk S-protein peptide library**

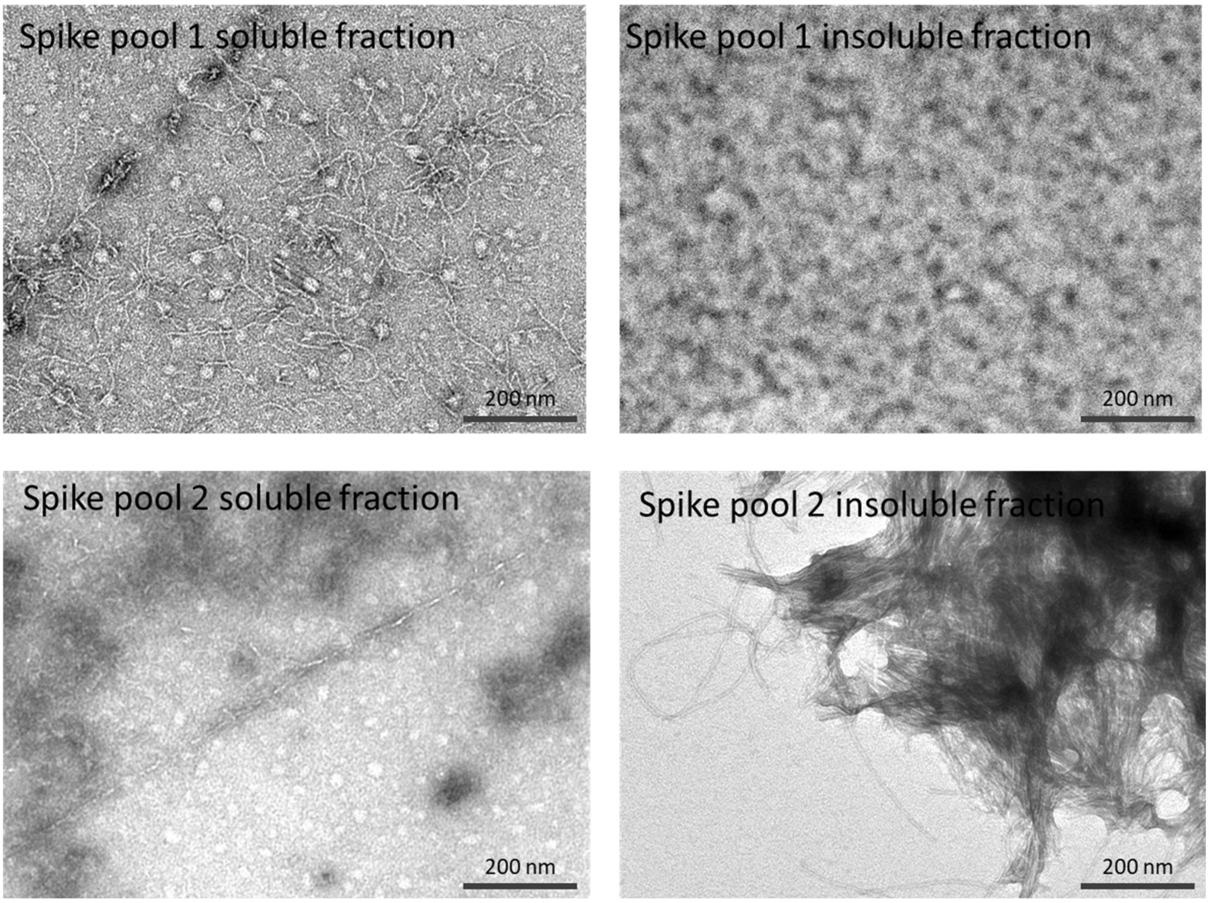


**Supporting Figure 1**. Negative stain TEM micrographs of **A.** Long single filament fibrils formed from PBS soluble fraction S-protein peptide library pool 1. **B.** Amorphous aggregates from PBS insoluble fraction S-protein peptide library pool 1. **C.** Short fibrils and amorphous aggregates formed from PBS soluble fraction S-protein peptide library pool 2 **B.** Heavily clustered long fibrils from PBS insoluble fraction S-protein peptide library pool 2.

**Supporting Figure 2: MALDI-ToF mass spectrometry spectra of synthesized selected peptides**
The mass indicated by the major single positive charged peak as well as the theoretical mass for each peptide is indicated in each spectrum

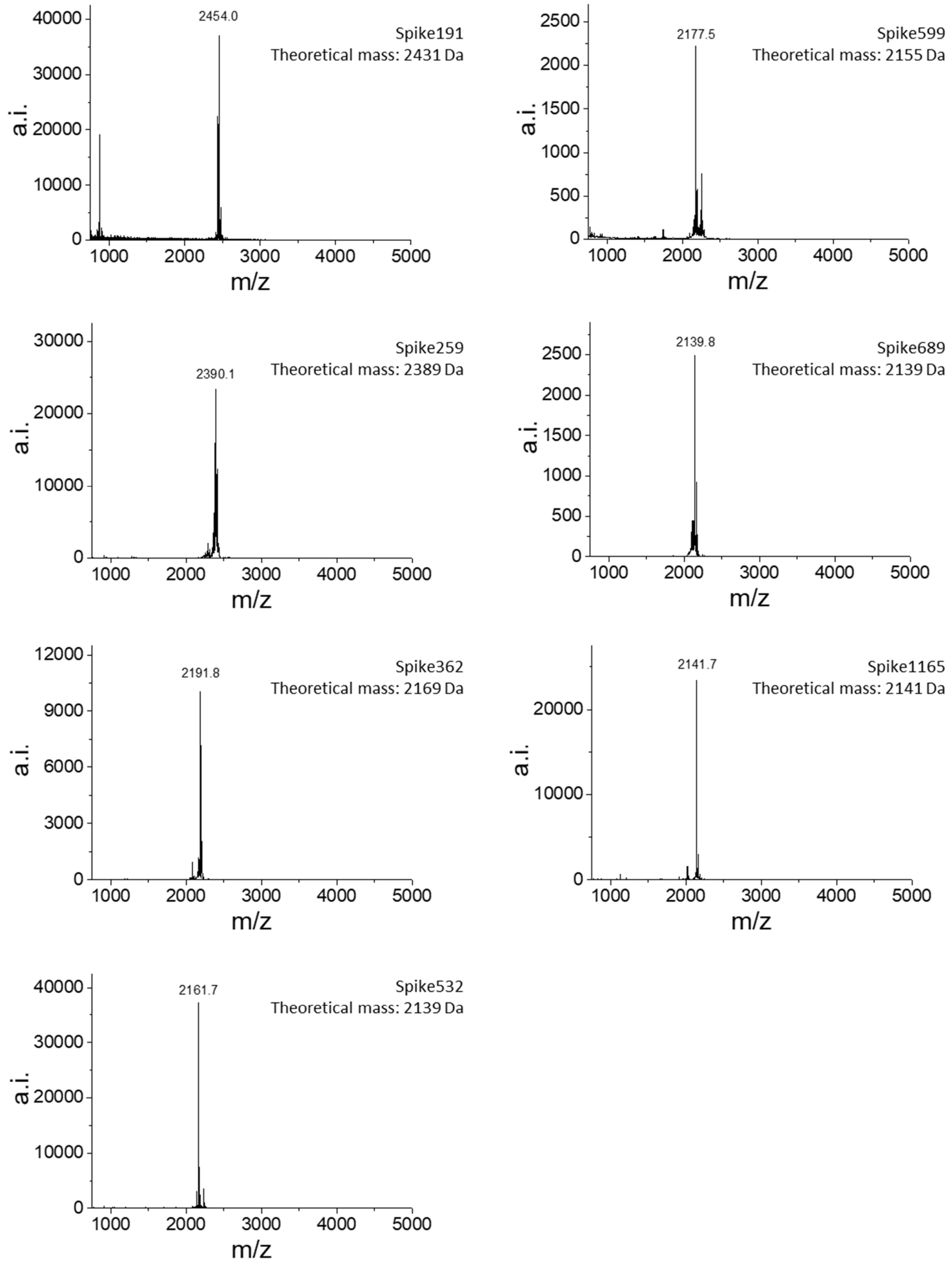


**Supporting Figure 2**. MALDI-ToF spectra of synthetic peptides dissolved in PBS buffer (10% HFIP) at 0.1 mg/ml to verify sequence and purity. Spike191, Spike362, Spike532, Spike599 show +22 Da due to $Na^+$ complex formation.

**Supporting Figure 3: Thermal denaturation**

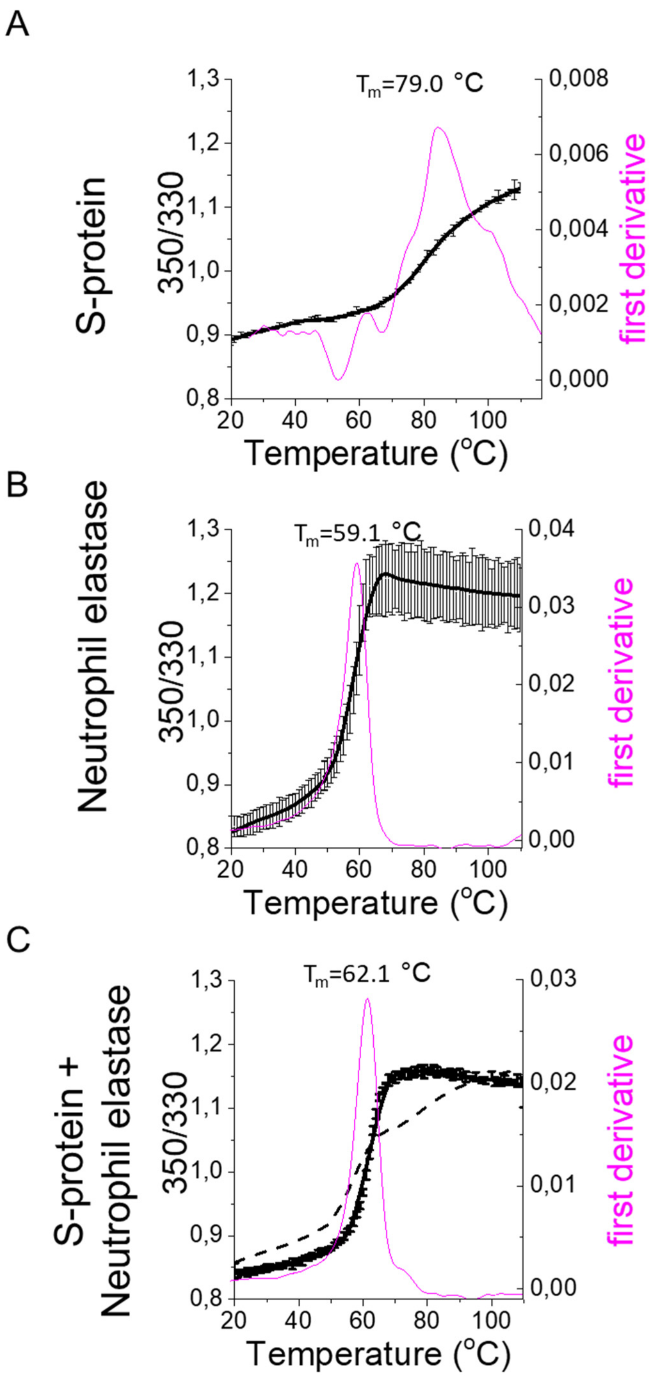


**Supporting figure 3:** Thermal denaturation by differential scanning fluorimetry (DSF) at pH 8.4 of A) S-protein B) Neutrophil elastase C) co-incubation of S-protein and Neutrophil elastase. Black curves show increase in tryptophan exposure to polar environment by emission ratio 350nm/330nm. Data is the average of two separate experiments with standard deviations. The dashed line in (C) represents the summed signals from A and B which does not fit the experimental data of the co-incubated proteins indicating S-protein cleavage by elastase. Magenta curve shows the first derivative of the thermal denaturation curve, accentuating the infliction points during denaturation. S-protein (A) in particular, displays a complex denaturation transition, dictated by the dissociation of the trimer as well as unfolding of the different subunits within each monomer.

**Supporting Figure 4: Raw transmission electron micrographs of full-length SARS-CoV-2 S-protein, Neutrophil Elastase and S-protein+Elastase incubated at 37 °C for 24h**

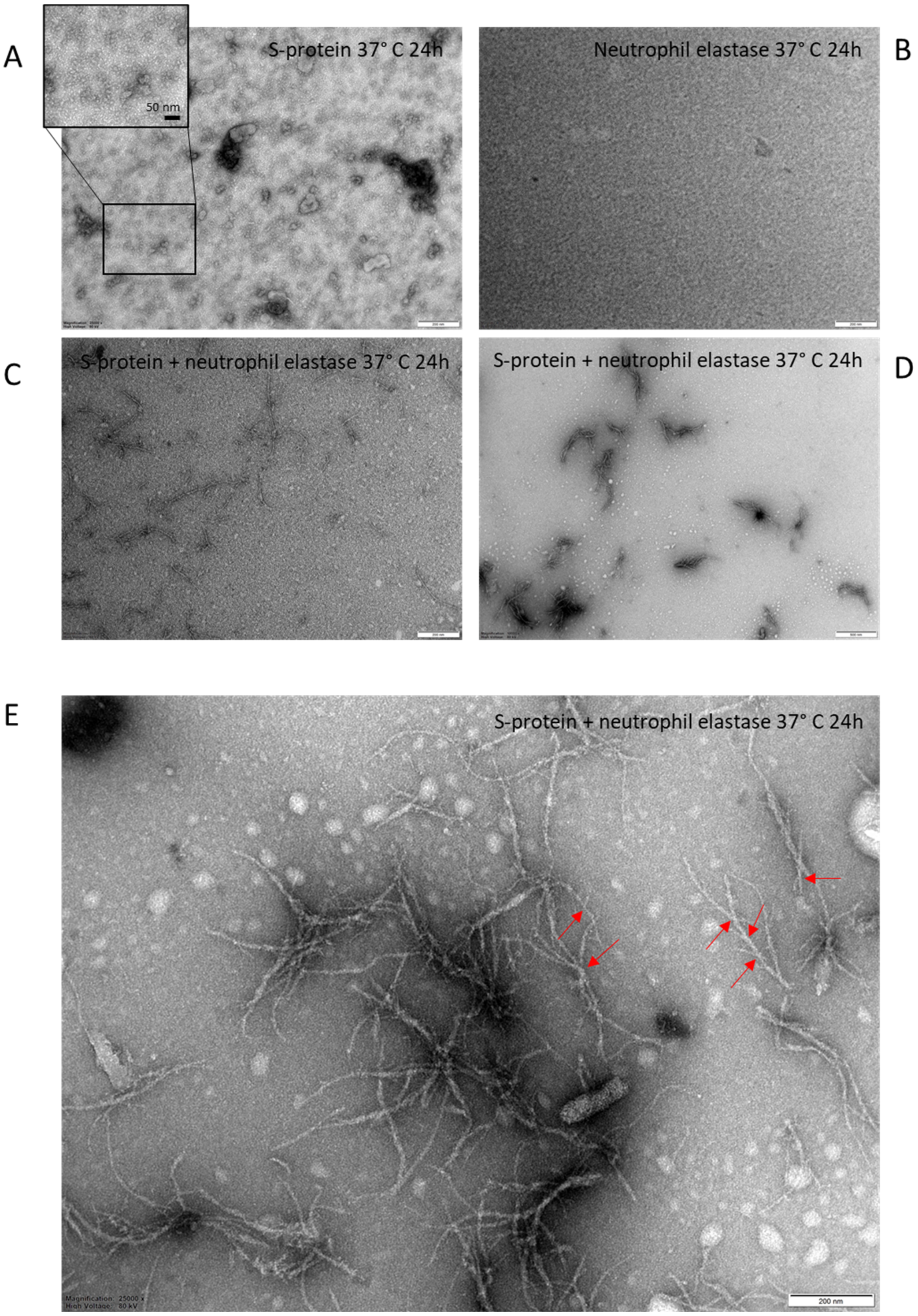


**Supporting Figure 4.** Negative stain TEM micrographs of **A.** S-protein, inset highlights the trimeric spike protein as displayed in [2] **B.** neutrophil elastase and **C-E**. S-Protein + elastase co-incubated in vitro at 37 °C for 24 h. S-protein trimer structures are visible in **A.** Fibrils were only formed in the co-incubated samples showing single filament fibrils in **C** and clusters of amyloid-like fibrils in **D.** The unusual fibril morphology with evident branching is clearly visible in the high magnification image in **E** and are highlighted with red arrows.

**Supporting Figure 5: WALTZ amyloidogenic sequence predictions of S-protein sequences from seven corona viruses known to infect humans**

A

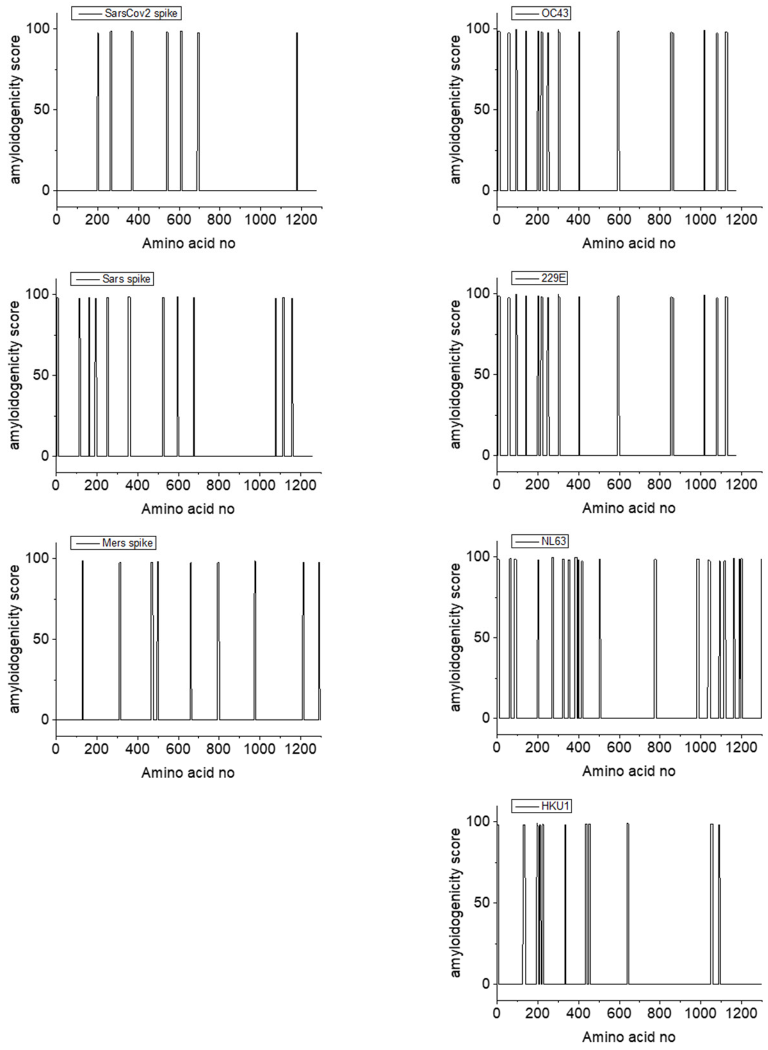


B

```
MERS              EDEILEWFGITQTAQGVHLFSSRYVD----LYGG--------------N-MFQFATLPVYD
SARS              SLDVSEKSG---NFKHLREFVFKNKDGFLYVYKGYQPIDVVRDLPSGFNTLKPIFKLPLGI
SARSCOV2          LMDLEGKQG---NFKNLREFVFKNIDGYFKIYSKHTPINLVRDLPQGFSALEPLVDLPIGI
                    ::    *   . : :: *  :  *    :*                . :  :  **:
                174                                                           232

OC43              TYDVNAT------YLYFHFYQEGGTFYAYFTDTG------FVTKF-----LFNVYLGM
229E              NGTNTSH------SVCNGCVGHSENVFAVESGGY------IPSNF-----AFNNWFLL
NL63              NGRIVNY------TVCDDCNGYTDNIFSVQQDGR------IPNGF-----SFNNWFLL
HKU1              FTYNVSTD------WLYFFYQERGTFYAYYADSG------MPTTF-----LFSLYLGT
SARSCOV2          LMDLEGKQGNFKNLREFVFKNIDGYFKIYSKHTPINLVRDLPQGFSALEPLVDLPIGI
                                           .              :   *      ..  :
                174                                                        232
```

**Supporting Figure 5 A)** WALTZ prediction of amyloidogenic sequences in Spike proteins from several corona viruses known to infect humans **B)** Sequence alignments of spike191 and flanking amino acids from the three corona viruses causing severe disease. MERS S-protein does not contain the amyloidogenic sequence, SARS S-protein contains a similar amyloidogenic sequence as SARS-CoV-2 S-protein in this part of the amino acid sequence. SARS-CoV-2 S-protein segment 193-212, highlighted in red was a peptide predicted for elastase cleavage.

**Supporting Figure 6: Fluorescence micrographs and spectra of Spike191 fibrils stained with fluorescent analogs of amyloid PET tracers**

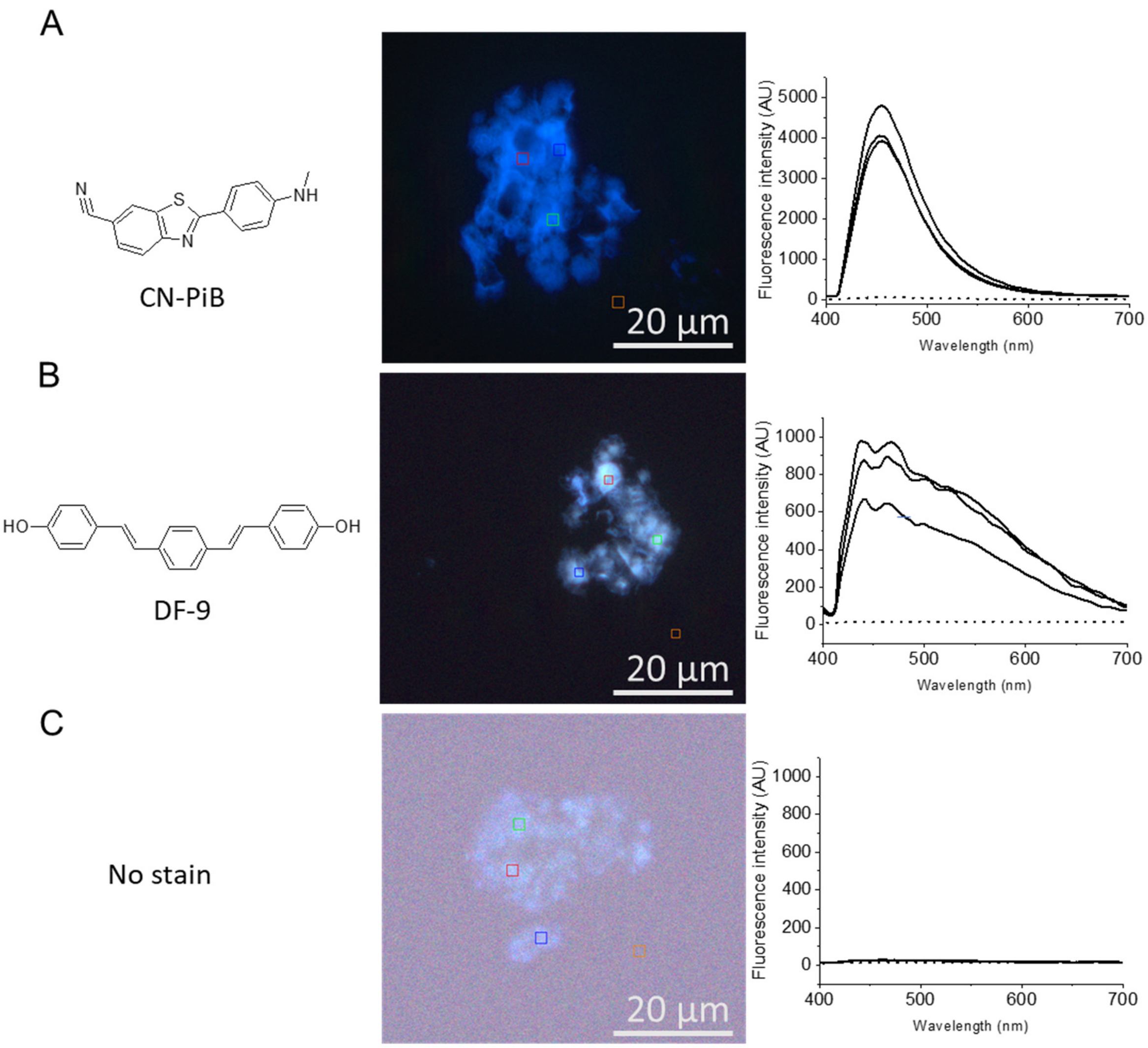


**Supporting Figure 6.** Fibrils of Spike191 stained with ligands CN-PiB (A) and DF-9 (B), fluorescent analogues of the PET tracers $^{11}$C-PiB and $^{18}$F-Florbetaben respectively, imaged with fluorescence hyperspectral microscopy. Both ligands show intense staining of fibrils with the expected emission spectra bound to amyloid fibrils. Spectra from three regions of interest (ROI) are depicted to the right of the micrographs, marked with colored squares in the micrographs. The brown colored ROIs are background signals outside of the aggregates and are as expected at very low intensity in the spectral graphs (dotted line). Autofluorescence (C) from Spike191 fibrils at identical microscope settings shown in C is negligible as evident from the poor image contrast and intensity of the spectra.

### Supporting Table 1 (STable 1)

| Accession | Description | Score A2 | Coverage A2 | # Peptides A2 | # PSM A2 | Score B2 | Coverage B2 | # Peptides B2 | # PSM B2 | # AAs | MW [kDa] | calc. pI |
|---|---|---|---|---|---|---|---|---|---|---|---|---|
| P0DTC2 | Spike glycoprotein OS=Severe acute respiratory syndrome coronavi | 254,74 | 26,94 | 66 | 87 | 162,40 | 17,99 | 39 | 60 | 1273 | 141,1 | 6,65 |

| position | Sequence | # PSMs | # Proteins | # Protein Groups | Protein Group Accessions | Modifications | MH+ [Da] | 1 min | XCorr 1min | 6 h | XCorr 6 h | # Missed Cleavages | missed clevage unique 6h | missed clevage 1 min |
|---|---|---|---|---|---|---|---|---|---|---|---|---|---|---|
| | LTESNKKFLPFQQFGRDIA | 1 | 1 | 1 | P0DTC2 | | 2239,19437 | | | High | 4,12 | 2 | 2 | 2 |
| | LTESNKKFLPFQQFGRDI | 2 | 1 | 1 | P0DTC2 | | 2168,15923 | | | High | 3,94 | 1 | 1 | 1 |
| | TVLPPLLTDEMI | 3 | 1 | 1 | P0DTC2 | | 1341,73052 | High | 2,36 | High | 3,90 | 3 | 3 | 3 |
| | AVEQDKNTQEV | 4 | 1 | 1 | P0DTC2 | | 1260,60564 | High | 2,45 | High | 3,83 | 2 | 2 | 2 |
| | AWNSNNLDSKV | 10 | 1 | 1 | P0DTC2 | | 1247,60503 | High | 3,06 | High | 3,74 | 2 | 2 | 2 |
| | RQIAPGQTGKI | 2 | 1 | 1 | P0DTC2 | | 1168,68328 | High | 2,69 | High | 3,40 | 1 | 1 | 1 |
| spike191 | FKNIDGYFKI | 4 | 1 | 1 | P0DTC2 | | 1244,66924 | | | High | 3,16 | 1 | 1 | 1 |
| | WNSNNLDSKV | 2 | 1 | 1 | P0DTC2 | | 1176,56926 | | | High | 3,07 | 1 | 1 | 1 |
| | IGKIQDSLSSTA | 2 | 1 | 1 | P0DTC2 | | 1219,65068 | High | 3,89 | High | 2,91 | 3 | 3 | 3 |
| | AKNLNESLI | 1 | 1 | 1 | P0DTC2 | | 1001,56664 | | | High | 2,90 | 3 | 3 | 3 |
| | SALGKLQDVV | 3 | 1 | 1 | P0DTC2 | | 1029,59734 | High | 2,53 | High | 2,86 | 4 | 4 | 4 |
| | IGKIQDSL | 1 | 1 | 1 | P0DTC2 | | 873,50768 | | | High | 2,80 | 2 | 2 | 2 |
| | RQIAPGQTGKIA | 1 | 1 | 1 | P0DTC2 | | 1239,72270 | | | High | 2,80 | 2 | 2 | 2 |
| | LPPLLTDEMI | 4 | 1 | 1 | P0DTC2 | M9(Oxidation) | 1157,61296 | | | High | 2,79 | 2 | 2 | 2 |
| | MSFPQSAPHGVV | 1 | 1 | 1 | P0DTC2 | | 1256,61540 | | | High | 2,78 | 1 | 1 | 1 |
| | KQIYKTPPI | 1 | 1 | 1 | P0DTC2 | | 1087,65569 | | | High | 2,75 | 1 | 1 | 1 |
| | VEQDKNTQEV | 1 | 1 | 1 | P0DTC2 | | 1189,57170 | | | High | 2,75 | 1 | 1 | 1 |
| | TQRNFYEPQII | 2 | 1 | 1 | P0DTC2 | | 1408,72661 | High | 2,16 | High | 2,75 | 1 | 1 | 1 |
| | YDPLQPELDSFKEEL | 1 | 1 | 1 | P0DTC2 | | 1822,87346 | | | High | 2,73 | 2 | 2 | 2 |
| | RDLPQGFSAL | 1 | 1 | 1 | P0DTC2 | | 1103,58953 | | | High | 2,64 | 1 | 1 | 1 |
| | GYLQPRTFLL | 2 | 1 | 1 | P0DTC2 | | 1207,68633 | High | 2,37 | High | 2,57 | 2 | 2 | 2 |
| | LYENQKLI | 2 | 1 | 1 | P0DTC2 | | 1020,57640 | High | 2,02 | High | 2,54 | 2 | 2 | 2 |
| | EDLLFNKV | 1 | 1 | 1 | P0DTC2 | | 977,53618 | | | High | 2,53 | 2 | 2 | 2 |
| | ANQFNSAI | 2 | 1 | 1 | P0DTC2 | | 864,42601 | High | 2,44 | High | 2,48 | 2 | 2 | 2 |
| | RDLPQGFSALEPL | 1 | 1 | 1 | P0DTC2 | | 1442,76299 | | | High | 2,46 | 2 | 2 | 2 |
| | SALGKLQDV | 2 | 1 | 1 | P0DTC2 | | 930,52971 | High | 2,38 | High | 2,40 | 3 | 3 | 3 |
| | GGNYNYL | 1 | 1 | 1 | P0DTC2 | | 800,36003 | | | High | 2,35 | 0 | 0 | 0 |
| | LGKLQDV | 1 | 1 | 1 | P0DTC2 | | 772,45989 | | | High | 2,33 | 2 | 2 | 2 |
| | GKIQDSLSSTA | 2 | 1 | 1 | P0DTC2 | | 1106,56877 | High | 3,22 | High | 2,31 | 2 | 2 | 2 |
| | GKIQDSL | 1 | 1 | 1 | P0DTC2 | | 760,42369 | | | High | 2,30 | 1 | 1 | 1 |
| | YKTPPI | 2 | 1 | 1 | P0DTC2 | | 718,41649 | | | High | 2,18 | 0 | 0 | 0 |
| | TLADAGFI | 1 | 1 | 1 | P0DTC2 | | 807,42784 | | | High | 2,17 | 3 | 3 | 3 |
| | MSFPQSAPHGV | 1 | 1 | 1 | P0DTC2 | | 1157,54509 | | | High | 2,16 | 0 | 0 | 0 |
| | GYQPYRVV | 2 | 1 | 1 | P0DTC2 | | 981,51518 | | | High | 2,10 | 1 | 1 | 1 |
| | AWNSNNL | 1 | 1 | 1 | P0DTC2 | | 818,38158 | | | High | 2,09 | 1 | 1 | 1 |
| | VNIQKEI | 2 | 1 | 1 | P0DTC2 | | 843,49285 | High | 2,24 | High | 2,07 | 2 | 2 | 2 |
| | APGQTGKIA | 1 | 1 | 1 | P0DTC2 | | 842,47209 | | | High | 2,05 | 1 | 1 | 1 |
| | DGYFKI | 1 | 1 | 1 | P0DTC2 | | 742,37956 | | | High | 2,05 | 0 | 0 | 0 |
| | GYLQPRTFL | 1 | 1 | 1 | P0DTC2 | | 1094,60137 | | | High | 2,03 | 1 | 1 | 1 |
| | RAAEIRA | 1 | 1 | 1 | P0DTC2 | | 786,45867 | High | 2,54 | | | 3 | **1,307692308** | 3 |
| | RAAEIRASA | 1 | 1 | 1 | P0DTC2 | | 944,52818 | High | 2,06 | | | 4 | **0,837578929** | 4 |
| | FAQVKQI | 1 | 1 | 1 | P0DTC2 | | 833,48754 | High | 2,03 | | | 2 | | 2 |
| | AVEQDKNTQEVFA | 1 | 1 | 1 | P0DTC2 | | 1478,71086 | High | 2,47 | | | 3 | | 3 |
| | YSTGSNVFQTRA | 1 | 1 | 1 | P0DTC2 | | 1330,63835 | High | 3,09 | | | 1 | | 1 |
| | AYTMSLGAENSV | 3 | 1 | 1 | P0DTC2 | M4(Oxidation) | 1258,56145 | High | 3,85 | | | 3 | | 3 |
| | GKYEQYI | 1 | 1 | 1 | P0DTC2 | | 900,44597 | High | 2,07 | | | 0 | | 0 |
| | LYENQKLIA | 1 | 1 | 1 | P0DTC2 | | 1091,60954 | High | 2,14 | | | 3 | | 3 |
| | LTGIAVEQDKNTQEV | 1 | 1 | 1 | P0DTC2 | | 1644,84636 | High | 4,31 | | | 4 | | 4 |
| | KNLNESLI | 1 | 1 | 1 | P0DTC2 | | 930,52519 | High | 2,14 | | | 2 | | 2 |
| | KQLSSNFGAI | 1 | 1 | 1 | P0DTC2 | | 1064,56914 | High | 2,45 | | | 2 | | 2 |
| | EQDKNTQEVFAQV | 1 | 1 | 1 | P0DTC2 | | 1535,73357 | High | 2,85 | | | 2 | | 2 |
| spike1165 | SVVNIQKEI | 1 | 1 | 1 | P0DTC2 | | 1029,59465 | High | 2,35 | | | 3 | | 3 |
| | YFASTEKSNII | 1 | 1 | 1 | P0DTC2 | | 1272,63933 | High | 2,49 | | | 2 | | 2 |
| | IAYTMSLGAENSV | 2 | 1 | 1 | P0DTC2 | M5(Oxidation) | 1371,64519 | High | 3,24 | | | 4 | | 4 |
| | AVEQDKNTQEVFAQV | 1 | 1 | 1 | P0DTC2 | | 1705,83891 | High | 3,24 | | | 4 | | 4 |
| | HADQLTPTWRV | 1 | 1 | 1 | P0DTC2 | | 1323,67693 | High | 2,48 | | | 2 | | 2 |
| | NIQKEIDRLNEV | 1 | 1 | 1 | P0DTC2 | | 1470,79033 | High | 3,39 | | | 3 | | 3 |
| | NIQKEIDRLNEVA | 2 | 1 | 1 | P0DTC2 | | 1541,82659 | High | 3,53 | | | 4 | | 4 |
| | ANQFNSAIGKI | 1 | 1 | 1 | P0DTC2 | | 1162,62065 | High | 2,06 | | | 3 | | 3 |
| | KQLSSNFGAISSV | 1 | 1 | 1 | P0DTC2 | | 1337,70989 | High | 2,88 | | | 3 | | 3 |
| | NQNAQALNTLV | 1 | 1 | 1 | P0DTC2 | | 1185,62212 | High | 2,57 | | | 4 | | 4 |
| | LYENQKLIANQFNSA | 2 | 1 | 1 | P0DTC2 | | 1752,89311 | High | 3,56 | | | 4 | | 4 |
| | AYTMSLGAENSV | 1 | 1 | 1 | P0DTC2 | | 1242,56780 | High | 4,06 | | | 3 | | 3 |
| | VNQNAQALNTLV | 1 | 1 | 1 | P0DTC2 | | 1284,69158 | High | 3,14 | | | 5 | | 5 |
| | TQNVLYENQKLI | 1 | 1 | 1 | P0DTC2 | | 1462,79009 | High | 3,48 | | | 3 | | 3 |
| | YRFNGIGVTQNV | 1 | 1 | 1 | P0DTC2 | | 1367,70720 | High | 2,09 | | | 2 | | 2 |
| | SALGKLQDVVNQNA | 1 | 1 | 1 | P0DTC2 | | 1456,77556 | High | 3,22 | | | 5 | | 5 |
| | AIHADQLTPTWRV | 1 | 1 | 1 | P0DTC2 | | 1507,79851 | High | 3,41 | | | 4 | | 4 |
| | TGIAVEQDKNTQEVFAQV | 1 | 1 | 1 | P0DTC2 | | 1976,99748 | High | 2,61 | | | 5 | | 5 |
| | VNIQKEIDRLNEV | 1 | 1 | 1 | P0DTC2 | | 1569,87041 | High | 2,51 | | | 4 | | 4 |
| | IAYTMSLGAENSV | 1 | 1 | 1 | P0DTC2 | | 1355,65288 | High | 4,20 | | | 4 | | 4 |
| | TWFHAI | 1 | 1 | 1 | P0DTC2 | | 774,39354 | High | 2,01 | | | 1 | | 1 |
| | HADQLTPTWRVYSTGSNVFQTRA | 2 | 1 | 1 | P0DTC2 | | 2635,29233 | High | 3,51 | | | 4 | | 4 |
| | SVTTEILPV | 1 | 1 | 1 | P0DTC2 | | 958,54686 | High | 2,13 | | | 2 | | 2 |
| | DRLNEVAKNLNESLI | 1 | 1 | 1 | P0DTC2 | | 1727,93035 | High | 3,03 | | | 5 | | 5 |
| | LYENQKLIANQFNSAI | 2 | 1 | 1 | P0DTC2 | | 1865,97246 | High | 3,78 | | | 5 | | 5 |
| | ALQIPFAMQMA | 2 | 1 | 1 | P0DTC2 | M8(Oxidation); M10(Oxidation) | 1252,60222 | High | 2,46 | | | 3 | | 3 |
| | TQNVLYENQKLIANQFNSA | 2 | 1 | 1 | P0DTC2 | | 2195,10723 | High | 4,63 | | | 5 | | 5 |
| | SALGKLQDVVNQNAQAL | 1 | 1 | 1 | P0DTC2 | | 1768,96355 | High | 2,75 | | | 7 | | 7 |
| | ANQFNSAIGKIQDSLSSTA | 1 | 1 | 1 | P0DTC2 | | 1951,97209 | High | 5,04 | | | 5 | | 5 |
| | NQFNSAIGKIQDSLSSTA | 1 | 1 | 1 | P0DTC2 | | 1880,93511 | High | 4,52 | | | 4 | | 4 |
| | LYENQKLIANQFNSAIGKI | 1 | 1 | 1 | P0DTC2 | | 2164,17386 | High | 3,18 | | | 6 | | 6 |
| spike689 | ASQSIIAYTMSLGAENSV | 1 | 1 | 1 | P0DTC2 | M10(Oxidation) | 1857,88970 | High | 5,21 | | | 6 | | 6 |
| | NFNFNGLTGTGV | 1 | 1 | 1 | P0DTC2 | | 1240,59905 | High | 2,53 | | | 1 | | 1 |
| | LTESNKKFLPFQQFGRDIADTTDAV | 2 | 1 | 1 | P0DTC2 | | 2841,43906 | High | 3,19 | | | 4 | | 4 |
| | VLSFELLHAPA | 1 | 1 | 1 | P0DTC2 | | 1196,66826 | High | 3,78 | | | 4 | | 4 |
| | DLQELGKYEQYI | 4 | 1 | 1 | P0DTC2 | | 1498,73955 | High | 3,89 | | | 2 | | 2 |
| | IANQFNSAIGKIQDSLSSTA | 1 | 1 | 1 | P0DTC2 | | 2065,05889 | High | 3,70 | | | 6 | | 6 |
| | ALQIPFAMQMA | 2 | 1 | 1 | P0DTC2 | M8(Oxidation) | 1236,61113 | High | 2,64 | | | 3 | | 3 |
| | TQNVLYENQKLIANQFNSAI | 1 | 1 | 1 | P0DTC2 | | 2308,19199 | High | 4,04 | | | 6 | | 6 |
| | TGRLQSLQTYVTQQLI | 1 | 1 | 1 | P0DTC2 | | 1849,01763 | High | 3,16 | | | 4 | | 4 |
| | LPPLLTDEMIA | 1 | 1 | 1 | P0DTC2 | M9(Oxidation) | 1228,64922 | High | 2,21 | | | 3 | | 3 |
| | NIQKEIDRLNEVAKNLNESLI | 1 | 1 | 1 | P0DTC2 | | 2453,33805 | High | 4,06 | | | 7 | | 7 |
| | TVLPPLLTDEMI | 1 | 1 | 1 | P0DTC2 | M11(Oxidation) | 1357,72563 | High | 3,08 | | | 3 | | 3 |
| | ASQSIIAYTMSLGAENSV | 1 | 1 | 1 | P0DTC2 | | 1841,90117 | High | 2,75 | | | 6 | | 6 |
| | ALQIPFAMQMA | 1 | 1 | 1 | P0DTC2 | | 1220,61687 | High | 3,04 | | | 3 | | 3 |
| | LPPLLTDEMI | 1 | 1 | 1 | P0DTC2 | | 1141,61711 | High | 2,12 | | | 2 | | 2 |
| spike259 | YYVGYLQPRTFLL | 1 | 1 | 1 | P0DTC2 | | 1632,87432 | High | 2,71 | | | 3 | | 3 |
| | | | | | | | | | | | | | | **3,319444444** |
| | | | | | | | | | | | | | | **1,499543488** |

### Supporting Table 2 (STable 2)

| Checked | Confidence | Sequence | # PSMs | # Missed Cleavages | Theo. MH+ [Da] | Abundance: 1 min: Sample | Abundance: 6 h: Sample | Abundance: No elastase: Sample | Abundances Count: 1 min: Sample | Abundance s Count: 6 h: Sample | Abundances Count: No elastase: Sample | Quan Info | Found in Sample: [S1] F1: Sample | Found in Sample: [S2] F2: Sample | Found in Sample: [S3] F3: Sample | Confidence (by Search Engine): Sequest HT | XCorr (by Search Engine): Sequest HT | Top Apex RT [min] |
|---|---|---|---|---|---|---|---|---|---|---|---|---|---|---|---|---|---|---|
| | High | TVLPPLLTDEMI | 4 | 3 | 1341,73342 | 6950,01709 | 44997,10156 | | 1 | 1 | | | Low | High | Not Found | High | 3,6 | 66,62 |
| | High | RQIAPGQTGKI | 4 | 1 | 1168,67968 | 53566,72461 | 6286004,5 | 7056,859375 | 2 | 2 | 1 | | High | High | Peak Found | High | 3,51 | 23,69 |
| | High | LYENQKLIANQFNSAI | 3 | 5 | 1865,97559 | 190799,3066 | | | 3 | | | | High | Not Found | Not Found | High | 3,07 | 49,52 |
| | High | AYTMSLGAENSV | 3 | 3 | 1258,56199 | 752863,4844 | | | 3 | | | | High | Not Found | Not Found | High | 3,05 | 34,49 |
| | High | IGIVNNTVYDPLQPELDSFKEELDKYFKNHTSPDV | 1 | 7 | 4065,01277 | 11067,69824 | | | 1 | | | | High | Not Found | Not Found | High | 2,92 | 50,73 |
| | High | GYLQPRTFLL | 7 | 2 | 1207,68337 | 155320,3984 | 801219,2461 | 235345,2578 | 3 | 3 | 2 | | Low | High | Low | High | 2,88 | 54,65 |
| | High | ASQSIIAYTMSLGAENSV | 1 | 6 | 1857,88987 | 25198,77539 | | | 1 | | | | High | Not Found | Not Found | High | 2,77 | 54 |
| | High | LTGIAVEQDKNTQEV | 1 | 4 | 1644,84391 | 15909,20703 | | | 1 | | | | High | Not Found | Not Found | High | 2,72 | 34,71 |
| | High | AIHADQLTPTWRV | 2 | 4 | 1507,80159 | 323107,3203 | | | 2 | | | | High | Not Found | Not Found | High | 2,7 | 44,99 |
| | High | IGKIQDSLSSTA | 4 | 3 | 1219,65286 | 414803,4063 | 1016118,102 | 5343,184082 | 1 | 3 | 1 | | High | High | Peak Found | High | 2,68 | 31,78 |
| | High | TQNVLYENQKLI | 1 | 3 | 1462,79002 | 96766,64063 | | | 1 | | | | High | Not Found | Not Found | High | 2,62 | 44,05 |
| | High | FKNIDGYFKI | 4 | 1 | 1244,66739 | 34703,43945 | 1001409,891 | | 2 | 4 | | | Peak Found | High | Not Found | High | 2,6 | 48,81 |
| | High | IDDHFLFDKPVSPLLLASGMARDWPDA | 1 | 8 | 3026,50805 | 48199,02734 | 145140,875 | | 1 | 1 | | | High | Peak Found | Not Found | High | 2,59 | 35,4 |
| | High | RQIAPGQTGKIA | 7 | 2 | 1239,7168 | 22276,9104 | 1413105,562 | 22445,10986 | 2 | 4 | 3 | | Low | High | Low | High | 2,58 | 22,86 |
| | High | IGKIQDSL | 1 | 2 | 873,50401 | | 1974504,875 | | | 1 | | | Not Found | High | Not Found | High | 2,57 | 32,87 |
| | High | NDVTDEMVRLNWLTEFMPLPTIKHFIRTPDDAWLL | 1 | 8 | 4243,13548 | 13560,90137 | | | 1 | | | | High | Not Found | Not Found | High | 2,56 | 43,97 |
| | High | QIPFAMQMAYRFNGIGVTQNVLYENQKL | 1 | 6 | 3305,64456 | | 90251,64844 | | | 1 | | | Not Found | High | Not Found | High | 2,5 | 27,23 |
| | High | GWARGQPAAAPQPGLVPPARRHYSEAAADREDDPNFFKMV | 6 | 8 | 4376,15301 | 30210,33203 | 57305,63281 | 15917,47461 | 1 | 1 | 1 | | High | Low | Peak Found | High | 2,48 | 53,94 |
| | High | TQNVLYENQKLIANQFNSAI | 4 | 6 | 2308,19319 | 32521,11523 | | | 1 | | | | High | Not Found | Not Found | High | 2,48 | 56,05 |
| | High | WNSNNLDSKV | 2 | 1 | 1176,56438 | | 4565650 | | | 1 | | | Not Found | High | Not Found | High | 2,44 | 33,65 |
| | High | AVEQDKNTQEV | 3 | 2 | 1260,60664 | 916284,9375 | 12008247 | | 1 | 1 | | | Low | High | Not Found | High | 2,41 | 18,7 |
| | High | DLQELGKYEQYI | 4 | 2 | 1498,7424 | 10457131,2 | | | 3 | | | | High | Not Found | Not Found | High | 2,3 | 55,33 |
| | High | IAYTMSLGAENSV | 3 | 4 | 1355,65114 | 124841,1641 | 7674521,738 | | 1 | 3 | | | High | Low | Not Found | High | 2,3 | 26,87 |
| | High | KQIYKTPPI | 3 | 1 | 1087,65101 | 108246,4219 | 2555684,289 | | 1 | 3 | | | Low | High | Not Found | High | 2,29 | 29,87 |
| | High | LYENQKLI | 4 | 2 | 1020,57242 | 864230 | 8517863,15 | | 1 | 2 | | | Low | High | Not Found | High | 2,27 | 35,77 |
| | High | LPPLLTDEMI | 6 | 2 | 1157,61224 | 18278,06641 | 203331,8286 | | 1 | 4 | | | Low | High | Not Found | High | 2,2 | 55,59 |
| | High | NQFNSAIGKIQDSLSSTA | 1 | 4 | 1880,93485 | 139279,9375 | | | 1 | | | | High | Not Found | Not Found | High | 2,17 | 53,29 |
| | High | GKIQDSLSSTA | 3 | 2 | 1106,56879 | 260163,7344 | 190775,25 | | 1 | 1 | | | High | Low | Low | High | 2,17 | 27,62 |
| | High | SALGKLQDVV | 6 | 4 | 1029,59389 | 339548,1563 | 7492608,186 | 53436,39063 | 2 | 2 | 1 | | Low | High | Low | High | 2,16 | 43,53 |
| | High | DDHFLFDKPV | 2 | 1 | 1232,59461 | | 73823,06055 | | | 2 | | | Not Found | High | Not Found | High | 2,15 | 44,77 |
| | High | AWNSNNLDSKV | 10 | 2 | 1247,60149 | 42988,97656 | 15111289,61 | 3520,373291 | 1 | 6 | 1 | | Low | High | Peak Found | High | 2,13 | 35,74 |
| | High | GIGTGEASLGGGGGGKV | 2 | 3 | 1373,70193 | 12548,38965 | 45165,05469 | | 1 | 1 | | | Low | High | Not Found | High | 2,13 | 27,59 |
| | High | ANQFNSAIGKIQDSLSSTA | 1 | 5 | 1951,97196 | 617012 | | | 1 | | | | High | Not Found | Not Found | High | 2,12 | 53,36 |
| | High | VLSFELLHAPA | 1 | 4 | 1196,66739 | 46165,73047 | 19356,16211 | | 1 | 1 | | | High | Peak Found | Not Found | High | 2,11 | 54,43 |
| | High | SALGKLQDV | 3 | 3 | 930,52547 | 129661,2891 | 4433163,5 | | 1 | 1 | | | High | Low | Not Found | High | 2,09 | 36,19 |
| | High | LGKLQDV | 2 | 2 | 772,45633 | | 467627,4727 | | | 2 | | | Not Found | High | Not Found | High | 2,09 | 28,95 |
| | High | AKNLNESLI | 18 | 3 | 1001,56259 | 9290699,796 | 13419974,1 | 8732547,089 | 10 | 7 | 8 | | Low | High | Low | High | 2,09 | 47,44 |
| | High | FAQVKQI | 3 | 2 | 833,48796 | 187353,4688 | | | 1 | | | | High | Not Found | Not Found | High | 2,09 | 30,38 |
| | High | AYTMSLGAENSV | 1 | 3 | 1242,56708 | 1436493,875 | | | 1 | | | | High | Not Found | Not Found | High | 2,03 | 43,07 |
| | High | GGNYNYL | 2 | 0 | 800,35734 | 5431,15625 | 1148047,973 | | 1 | 2 | | | Peak Found | High | Not Found | High | 2,01 | 37,5 |
| | High | KQLSSNFGAISSV | 3 | 3 | 1337,70596 | 96529,61719 | 98255,95313 | | 1 | 1 | | | High | Low | Low | High | 2,01 | 18,55 |
| | High | LPPLLTDEMI | 1 | 2 | 1141,61732 | 27614,21094 | 290739,2813 | 3666,937012 | 1 | 1 | 1 | | High | Peak Found | Peak Found | High | 1,99 | 61,9 |
| | High | NQNAQALNTLV | 1 | 4 | 1185,62223 | 98478,05469 | | | 1 | | | | High | Not Found | Not Found | High | 1,99 | 42,54 |
